# Demixer: A probabilistic generative model to delineate different strains of a microbial species in a mixed infection sample

**DOI:** 10.1101/2024.04.11.589150

**Authors:** VP Brintha, Manikandan Narayanan

## Abstract

**Motivation:** Multi-drug resistant or hetero-resistant Tuberculosis (TB) hinders the successful treatment of TB. Hetero-resistant TB occurs when multiple strains of the TB-causing bacterium with varying degrees of drug susceptibility are present in an individual. Existing studies predicting the proportion and identity of strains in a mixed infection sample rely on a reference database of known strains. A main challenge then is to identify *de novo* strains not present in the reference database, while quantifying the proportion of known strains.

**Results:** We present Demixer, a probabilistic generative model that uses a combination of reference-based and reference-free techniques to delineate mixed infection strains in whole genome sequencing (WGS) data. Demixer extends a topic model widely used in text mining to represent known mutations and discover novel ones. Parallelization and other heuristics enabled Demixer to process large datasets like CRyPTIC (Comprehensive Resistance Prediction for Tuberculosis: an International Consortium). In both synthetic and experimental benchmark datasets, our proposed method precisely detected the identity (e.g., 91.67% accuracy on the experimental *in vitro* dataset) as well as the proportions of the mixed strains. In real-world applications, Demixer revealed novel high confidence mixed infections (101 out of 1,963 Malawi samples analyzed), and new insights into the global frequency of mixed infection (2% at the most stringent threshold in the CRyPTIC dataset) and its significant association to drug resistance. Our approach is generalizable and hence applicable to any bacterial and viral WGS data.

**Availability:** All code relevant to Demixer is available at https://github.com/BIRDSgroup/Demixer.

**Supplementary information:** The Supplemental Data/Result Files related to Demixer are available at this link: https://drive.google.com/drive/folders/13WFACrn2EpeVTO7533-YwlAGjgF4UH3k?usp=drive_link.

## 1. Introduction

Tuberculosis (TB), a contagious disease caused by the bacterium *Mycobacterium tuberculosis (M. tb)*, has been present for thousands of years. The treatment regimen for TB involves taking a combination of antibiotic drugs over a prolonged period. In recent times, treating TB has become increasingly difficult due to the evolving nature of the TB-causing bacterium. Specifically, the growing resistance of different strains of the TB-causing bacterium (referred to simply as strains of TB or TB strains hereafter) to commonly used drugs [1] has extended the treatment regimen from months to years [2]. For instance, Beijing and Euro-American strains of TB are known to be more drug-resistant than the other types, possibly due to their high prevalence [3]. TB infections caused by a single drug-sensitive strain are highly susceptible to first-line treatments, including isoniazid and rifampicin [1]. In contrast, mixed infections involving multiple strains of TB can complicate treatment and are more challenging to manage [4, 5]. Mixed TB infection in an individual can arise due to transmission of different strains to the individual (co-infection) or due to evolution of an existing strain during treatment [6]. Additionally, TB infections in individuals with poor treatment conditions or HIV-TB co-infection can accelerate the risk of acquiring mixed strains due to compromised immunity, and hence increase the severity of the disease [7].

Determining the identity (mutational profile) and the proportions of the individual strains in mixed infection TB samples could greatly improve the clinical decision-making process for TB treatment. There are two categories of techniques for identifying the mixture of bacterial strains in a sample: traditional and modern. Traditional approaches are molecular biology-based and relatively cost-effective when compared to modern techniques. Various versions of molecular subtyping techniques, such as Polymerase Chain Reaction (PCR), Restriction Fragment Length Polymorphism (RFLP), and Mycobacterial Interspersed Repetitive Unit–Variable Number Tandem Repeat (MIRU-VNTR), have been widely used to detect mixed infection strains. However, these traditional methods have limitations. For example, experiments have shown that MIRU-VNTR failed to detect drug-resistant strains in mixed infection cases, highlighting the need for more advanced detection methods [8]. It can be challenging to detect individual strains when closely related strains are mixed in a sample with a relatively low proportion of the minority strain [9]. Modern methods based on Whole Genome Sequencing (WGS), such as ones enabled by Illumina/Nanopore technologies, have the potential to replace existing techniques as they generate an enormous amount of high-resolution data. The WGS data, if properly analyzed, can reveal the identity and proportion of different strains with high sensitivity [10].

The WGS-based methods for quantifying strains can be classified into two main categories: reference-based and reference-free. TBProfiler [11] is a popular reference-based tool that uses a comprehensive database of reference lineages and sublineages to delineate the strains in a sample. QuantTB is a state-of-the-art reference-based method that uses a curated list of single nucleotide polymorphisms (SNPs) from genome sequences of select TB bacterial isolates to identify and quantify mixed infections [12]. This method also determines the type of antibiotic resistance using a set of manually selected SNPs related to drug resistance in *M. tb*. But, the effectiveness of QuantTB relies on the diversity of the reference database, which limits its ability to detect new strains in the absence of related strains in the database. MixInfect [13] is a reference-free approach that uses the ratio of heterozygous alleles to quantify the mixed infection proportions. Although this method is simple and does not require any overhead of keeping track of the new strains, it reveals only the strains’ proportions and not their identity (mutational profile), and also cannot robustly handle cases when strains are present in equal proportions. SplitStrains [14] is a new statistical-based method that aligns with MixInfect due to its non-reliance on the reference database. The procedure uses a likelihood ratio test to resolve the heterogeneity of the samples, followed by the assignment of reads to a strain using the Expectation-Maximization (EM) algorithm and Naive Bayes classifier, in order to finally report the strain proportions. But, SplitStrains like MixInfect cannot reveal the identity of the strains.

A major challenge therefore with existing WGS-based methods like QuantTB is their strict reliance on reference databases, and with reference-free approaches MixInfect or SplitStrains is their difficulty to infer the identity (mutational profile) of *de novo* strains. To address these challenges, we have introduced a hybrid (reference-plus-*de novo*) approach called Demixer to infer the individual strain proportions and their corresponding mutations present in a WGS dataset. Demixer makes these inferences using a probabilistic generative model that models the strain producing each WGS read of a sample as a latent (hidden or unknown) variable. In a bit more detail, Demixer model assumes that each sample’s WGS reads are generated via a process that first generates the proportions of different strains mixed in a sample [15], and then the reads corresponding to the mixed strains. By learning the parameters of this generative process, including those corresponding to the latent variable of each WGS read, we can accurately infer/predict the distribution of strains in a new sample. The Demixer model is an extension of a popular topic model called Latent Dirichlet Allocation (LDA) [16] in text mining. We have chosen to extend LDA due to its simplicity and interpretability compared to word-embedding-based models such as word2vec, LDA2vec, or DNABERT [17, 18, 19]. To better suit the unique characteristics of genomic data, we introduced specific modifications to the LDA model, referred to as SNP-LDA. The SNP-LDA model, along with preprocessing and postprocessing steps, collectively constitute our method, Demixer.

A main advantage of Demixer compared to other existing approaches is its ability to perform multi-sample analysis. This allows Demixer to crucially use the commonalities and differences among multiple samples to refine the mutational profile of a reference strain (for instance to augment known mutations of the strain in the reference database with additional mutations present in the samples), extract the mutational profile of a *de novo* strain present in multiple samples, and quantify their proportions. In tandem with this advancement, we have also implemented parallelization heuristics to reduce the execution time of Demixer and thereby enhance its scalability. We have shown Demixer to be effective in delineating the strains in a mixture in various synthetic and real-world datasets. Demixer for instance inferred the strain identities of mixed samples in an *in vitro* dataset with an accuracy of 91.67%, which is comparable to or better than existing approaches. In synthetic benchmarks at different coverage levels, Demixer estimated the strain proportions even at 20x coverage with 99% accuracy. In another synthetic dataset, Demixer estimated the proportions of *de novo* strains which were absent in the reference database (with mean relative error 0.061). On application to real-world Malawi dataset, Demixer detected 101 new (previously unreported) mixed infections with high confidence out of the 1,963 samples analyzed. Demixer when applied to CRyPTIC dataset revealed new insights into the the global prevalence of mixed infection and its association with drug resistance. Furthermore, the parameters of the model obtained by training on CRyPTIC dataset can be used to test new samples, making the method more applicable for real-world diagnosis.

## 2. Materials and methods

### 2.1. Overview of Demixer method

Our proposed Demixer’s main contribution, relative to published reference-based or reference-free approaches, is in enabling multi-sample hybrid (reference-plus-*de-novo*) analysis. This analysis allows Demixer to infer the identity and proportions of different reference as well as new strains within the samples. Our Demixer method uses the WGS reads of each sample and a database of mutations in reference strains as inputs to determine the composition of the reference and *de novo* strains present in the samples. This involves the following three steps (as also illustrated in Figure 1A).

**Fig. 1.**
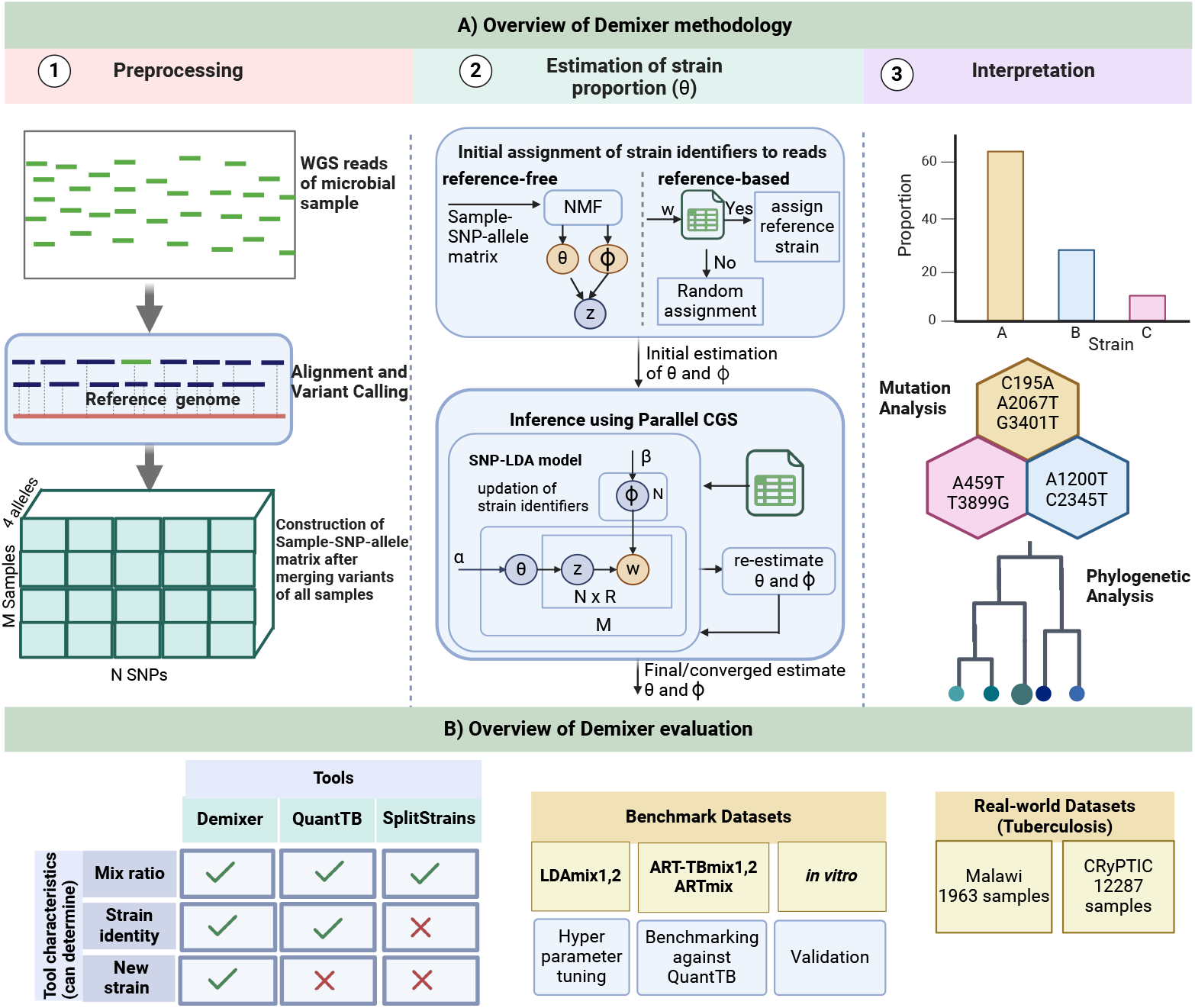
Our Demixer Overview: A) Illustrated here are the three steps of our Demixer method (preprocessing of WGS reads to generate the *M × N ×* 4 Sample-SNP-allele matrix, SNP-LDA model based inference of the Sample-Strain proportions ***θ*** from this matrix, and interpretation of the learnt mutations). Each SNP *n* of sample *m* is supported by *R* reads. Demixer includes a SNP-LDA model whose latent variable (strain identifier of each read) is initialized using NMF or known reference mutations. The matrix of parameters of our SNP-LDA model, ***θ*** and ***ϕ*** (Strain-SNP-allele distributions), are estimated using the parallelized Collapsed Gibbs Sampling (CGS) inference algorithm (refer Figure 2). B) Different synthetic/real-world datasets and methods used to assess the effectiveness of Demixer in estimating the strain identities and their proportions.

1. Preprocessing: Preprocessing the WGS reads of the input microbial samples involves performing read alignment and variant calling to generate the variants file (in .vcf format) for each sample, and merging them across all *M* samples to generate the Sample-SNP-allele matrix ***S*** of size *M × N ×* 4. Here, ***S***_*i,j,v*_ refers to the number of reads supporting the allele *v* of SNP *j* in sample *i* (with each SNP out of the *N* total SNPs taking one of the four alleles: A, C, G, or T).
2. Estimation of strain proportion: Demixer comprises a probabilistic generative model SNP-LDA, which extends the popular topic model LDA, to model the generative process of the Sample-SNP-allele matrix (as illustrated in Supplementary Figure S1 and Figure 1A.2). Demixer then estimates the parameters of this generative process (including the strain composition of samples and the distribution of mutations in strains) from the input Sample-SNP-allele matrix by performing inference on the SNP-LDA model. The inference can be guided by initializing the latent variable in the model representing the unknown strain of each WGS read using a database of unique mutations of reference strains or via Non-negative Matrix Factorization (NMF) [20].
3. Interpretation/Postprocessing: The third step involves the visualization and phylogeny/clustering-based interpretation of the SNP-allele distributions and proportions corresponding to different strains inferred by the model (after appropriate postprocessing steps).

One of the significant differences between our proposed Demixer and other methods is that the WGS samples are analyzed together instead of being processed separately. This integrated approach enables Demixer to harness any shared information across (similar) samples to better infer Sample-Strain compositions and Strain-SNP-allele distributions. Figure 1B shows the different datasets and state-of-the-art methods used to assess the performance of Demixer under diverse settings.

### 2.2. Background on LDA model

LDA, proposed by Blei et al. [16], is an unsupervised approach that is widely used in text mining to infer underlying topics in a corpus of documents. It belongs to the family of latent variable models, whose basic principle to model a given system is to employ a complex hidden mechanism (latent generative process) to link the unobserved (latent) variables to the observed variables in the system. Specifically, as shown next, LDA assumes that each document is generated using a mixture of *K* topics, and knowing the topic (latent variable *z*) assigned to each word *w* in the document could facilitate the estimation of document-topic distribution ***θ*** and word-topic distribution ***ϕ***.

- For each document *d*, generate ***θ***_*d*_ ∼ Dirichlet(*α*).
- For each topic *k*, generate ***ϕ***_*k*_ ∼ Dirichlet(*β*).
- To generate *n*-th word *w*_*n,d*_ in each document *d*:
  – Generate *z*_*n,d*_ ∼ Multinomial(***θ***_*d*_).
  – Generate *w*_*n,d*_ ∼ Multinomial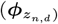.

Here, *α* and *β* are the hyperparameters of the LDA model; *z*_*n,d*_ is the latent variable capturing the topic assigned to word *w*_*n,d*_ at *n*^*th*^ position in document *d*. The generate term above refers to drawing/choosing/sampling a value for a random variable from the specified distribution. The model parameters are learnt from data *w* using variational or Gibb’s sampling based inference algorithms.

Recently, LDA has been used in bioinformatics applications involving large biological datasets analogous to LDA documents and words. Liu et al. [21] have categorized such applications into three types: clustering, classification, and feature extraction of biological data using the generative principle of LDA. Classic examples include gene sequence classification, metagenomic binning, and learning protein functions. Most applications use published LDA models directly to address specific challenges [21]. We had to modify and extend the LDA model to capture the reads assigned to the four alleles of a SNP (which has different semantics than words in a text); and we also propose a parallelization heuristic to speed up the CGS inference algorithm on our Demixer SNP-LDA model (Figure 1A).

### 2.3. Description of Demixer method

The three steps of our Demixer method overviewed briefly above, viz., Preprocessing, Estimation of strain proportion, and Interpretation (see also Figure 1A), are described in detail in this section.

#### 2.3.1. Preprocessing

Preprocessing involves different steps to process the FASTQ files containing raw reads of WGS samples to generate the Sample-SNP-allele matrix ***S***.

##### From WGS reads to genetic variants

A preprocessing pipeline for extracting the variants from WGS reads of samples has been developed. The quality of the raw reads is analyzed using the FastQC (version 0.11.9) [22] tool. Burrows-Wheeler Alignment (BWA) tool (version 0.7.17) [23] is used to map the raw reads in the sample against the H37Rv reference genome (NCBI Reference Sequence NC 000962.3). The output of BWA is subject to de-duplication and BAM file generation using the GATK tool (Genome Analysis Toolkit, version 4.1.8.0). From the resulting .bam file, FreeBayes (version 1.3.6) is used to call variants and generate the .vcf file. The .vcf files of individual samples are then merged using bcftools (version 1.14) [24]. A second round of variant calling is executed to call variants specifically at the positions in the merged .vcf file for each sample, which are then finally merged across all samples to get a single .vcf file. We then apply the Minor Allele Frequency (MAF) cutoff to remove the rare SNPs. As per standard practice, variants from PE/PPE gene regions are removed, and only SNPs from the coding regions are extracted for further analysis of the TB isolates [13].

The above pipeline is used to preprocess all the synthetic and benchmark datasets, but with slight modifications for the real-world datasets. We use dataset-specific MAF cutoff values to adapt to the different sizes of real-world datasets (Malawi - 0.002 and CRyPTIC - 0.0002; see Section 2.5 for details of these datasets). For the Malawi dataset, we applied the MAF filter both before (to speed up the process) and after the second variant calling on the merged vcf files. In the case of CRyPTIC, we used the processed .vcf files (with two rounds of variant calling) of individual samples obtained from their repository to get the final merged .vcf file and then applied the MAF cut-off.

##### Construction of Sample-SNP-allele matrix

The distinctive characteristics of different strains of a cellular organism are often related to specific genotypic features that serve as unique identifiers. One such genetic marker, SNPs within the DNA sequence, is widely used to distinguish the different strains reliably. A detailed analysis of the distribution of different alleles (A, C, G, T) of such SNPs present in a sample is required to detect mixed infection and delineate the proportion of the microbial strains in the sample [13]. This SNP-allele distribution is provided by the WGS reads obtained from a sample, with each read supporting a SNP-allele combination similar to a word in text LDA analysis [25]. The number of reads supporting each SNP-allele combination for all samples captures sufficient information for the model to learn the underlying strains, and we refer to this read counts matrix as the Sample-SNP-allele matrix ***S***. The ***S*** matrix is obtained from the single .vcf file mentioned above by including only the SNPs (i.e., by ignoring other variants like insertions and deletions). Very small entries in the ***S*** matrix (specifically any entry with less than 6 supporting reads) are made zero to mitigate noise from sequencing errors.

#### 2.3.2. Estimation of strain proportion

Demixer learns the strain proportion of samples via the SNP-LDA model. The notations, description, inference algorithm and heuristics adopted to improve the SNP-LDA model are explained in this section.

##### SNP-LDA model notations

The notations used to describe SNP-LDA model are given below. To simplify exposition, vectors, (two-dimensional) matrices, and three- or higher-dimensional tensors are all referred to as matrices in the text (with the matrix dimensions or sizes indicated as subscripts of the form *dim*_1_ *× dim*_2_ *×* …). We will need the indicator function 𝕝_condition_, which is defined as 1 if the condition (logical expression) evaluates to true, and 0 otherwise.

- Inputs:
  – *M* is the number of samples, *N* is the number of SNPs (obtained after preprocessing), and each read of a SNP supports one of the SNP’s 4 alleles: A, C, G, or T (also denoted 1, 2, 3, 4 respectively when used as index of a matrix).
  – ***S*** or ***S***_*M×N×*4_ - Sample-SNP-allele matrix of size *M × N ×* 4. Here, ***S***_*m,n,v*_ is the number of reads supporting allele *v* at *n*-th SNP in the *m*-th sample.
  – *R* - number of reads mapping to each SNP. This assumption of same number of reads per SNP simplifies exposition (however our model/implementation can easily handle different number of reads mapping to each SNP).
  – *w*_*m,n,r*_ - allele supported by the *r*-th read mapping to SNP *n* in sample *m*. Note that *S* can be constructed from all *w*_*m,n,r*_ observations as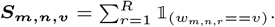.
  – *K* - Number of possible strains.
- Model parameters:
  – Sample-Strain distribution, ***θ***_*M×K*_.
  – Strain-SNP-allele distribution, ***ϕ***_*K×N×*4_
- Hyperparameters: *α* and *β*, which are the parameters of the Dirichlet distributions ***θ*** and ***ϕ*** respectively.
- *z*_*m,n,r*_ is the latent variable indicating the strain assigned to read *r* of *n*^*th*^ SNP in sample *m*.
- ***C***_*k,m,n,v*_ represents the number of reads of sample *m* at *n*^*th*^ SNP that supports allele *v* and is assigned to strain *k* (i.e., reads *r* for which *w*_*m,n,r*_ = *v* and *z*_*m,n,r*_ = *k*).
- Whenever an index is replaced by *∗* in any variable’s I:, notation above, we sum the variable over all possible values of the index. For instance, 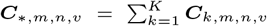***S***_*m,n,v*_ = ***C***_*∗,m,n,v*_, and ***S***_*m,n,∗*_ = *R*.

##### SNP-LDA model description

Each sample *m* is assumed to contain a mixture of *K* strains and is composed of a set of *N* SNPs, each quantified by *R* reads that reveal the composition of alleles at the SNP. The random variables of the model include the parameters ***θ***_*m*_ (distribution of strains in *m*-th sample) and ***ϕ***_*k,n*_ (distribution of the 4 alleles at SNP *n* for strain *k*), the latent variable *z*_*m,n,r*_ (strain assigned to the read denoted (*m, n, r*)), and the observed allele *w*_*m,n,r*_ ∈ {*A, C, G, T*}. The generative process of our SNP-LDA model follows:

- For each WGS sample *m*, generate ***θ***_*m*_ ∼ Dirichlet(*α*).
- For each strain *k* and SNP *n*, generate ***ϕ***_*k,n*_ ∼ Dirichlet(*β*_*n*_).
- To generate *r*-th read *w*_*m,n,r*_ at each SNP *n* in each sample *m*:
  – Generate *z*_*m,n,r*_ ∼ Multinomial(***θ***_*m*_).
  – Generate *w*_*m,n,r*_ ∼ Multinomial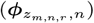.

Compared to the traditional LDA model’s generative process mentioned above, the main difference in the SNP-LDA model is to coherently model all the 4 alleles at each SNP position by using a ***ϕ*** matrix of size *K × N ×* 4. The advantage of this coherent modeling in SNP-LDA compared to LDA is also demonstrated empirically (see Results and Supplementary Section 1.6).

##### SNP-LDA model likelihood

The generative process of the SNP-LDA model (also captured in the DAG or Directed Acyclic Graph structure in Figure 1A.2) can be used to derive the likelihood of the model as shown below. First, let *w*_*m,·,·*_ refer to the set of alleles observed in all *R* reads of *N* SNPs of sample *m*, and let *z*_*m,·,·*_ be similarly defined as the set of the corresponding latent variables. Then, the joint distribution is given by:

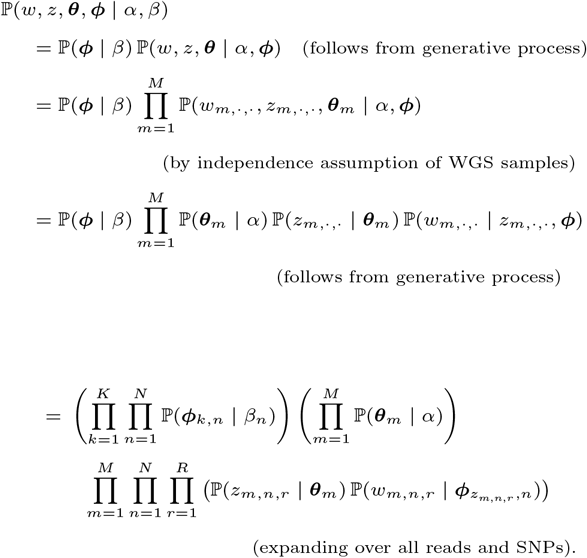

Note that ℙ above refers to the probability mass function for a discrete random variable and the probability density function for a continuous random variable.

##### SNP-LDA inference algorithm (Parameter estimation)

The posterior distribution of the latent variables given the observed data ℙ (***θ, ϕ***, *z* | *w, α, β*) is generally intractable, and Gibbs sampling offers one way to perform inference and parameter estimation by sampling from this posterior. The collapsed variant of a Gibbs Sampler (CGS) returns samples from the posterior ℙ (*z* | *w, α, β*), where the multinomial parameters ***θ*** and ***ϕ*** have been marginalized (collapsed) out.

Once the originating strain of each read given by *z* = {*z*_*m,n,r*_} is known (i.e., assigned via CGS samples), ***θ*** and ***ϕ*** can be easily estimated (see Supplementary Section 1.1).

One update step of CGS considers a specific read denoted (*m, n, r*), and assigns it to a strain (i.e., *z*_*m,n,r*_ is sampled/updated) given the strain assignment of all other reads (denoted by *z*_*−*(*m,n,r*)_). Let *w*_*m,n,r*_ = *v ∈ {A, C, G, T*}, and let *w*_*−*(*m,n,r*)_ denote the alleles at all other reads. If *z*_*m,n,r*_ = *k*^*′*^ before the updation step, define *C*^*−*(*m,n,r*)^ to be a copy of *C* but with one entry 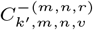 decremented by 1 (in order to remove the count contribution of the current read). Then, the conditional required for a CGS update step obtained from marginalizing the joint distribution above is given by Equation 1 and 2 below:

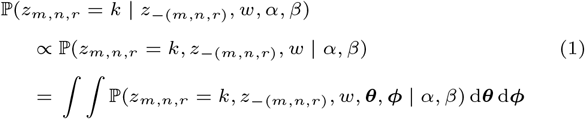

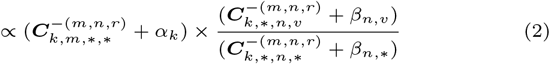

(recall *w*_*m,n,r*_ = *v*; see Supplementary Section 1.1 for derivation)

During initialization, i.e., before the Gibbs sampling iterations, each read is assigned to a random strain (which is picked uniformly at random from all the *K* strains; in other words, the strains of reads are initialized randomly). Then during one iteration/epoch of CGS, the strain assignment of all reads across all SNPs and samples are updated one at a time sequentially as per Equation 2. These CGS iterations are repeated 1000 times (as it leads to convergence in most of our experiments).

##### Heuristics to improve SNP-LDA model

We propose the following heuristics to improve strain identification and reduce the running time of the SNP-LDA model.

###### 1. Initialization heuristics

The SNP-LDA model may struggle to infer the proportions of minority strains that are mixed in very low frequencies in a sample. To address this issue and thereby enhance SNP-LDA model’s ability to precisely detect the Sample-Strain distribution, we propose two alternative methods to initialize the model parameters and latent variables.

- **NMF SNP-LDA:** This approach uses the output of NMF to initialize the strain of each read, instead of a random strain initialization described above. NMF is a widely used algorithm that factorizes a non-negative matrix into a product of two non-negative matrices to reveal certain (*K*^*′*^) underlying latent components as follows:

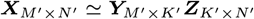

In our application, the observed data ***X*** is the Sample-SNP-allele matrix ***S***_*M×N×*4_ (collapsed into a 2D matrix of size *M ×* 4*N*), the inferred matrix ***Y*** represents the Sample-Strain distribution ***θ***_*M×K*_, and finally the other inferred matrix ***Z*** refers to the Strain-SNP-allele distribution ***ϕ***_*K×N×*4_ (collapsed into a *K ×* 4*N* sized 2D matrix). We used the NMF implementation from Python’s scikit-learn library [26], and chose Non-negative Double Singular Value Decomposition to initialize the NMF procedure.

As NMF is a simpler and efficient non-probabilistic approach, we apply it to factorize ***S*** into ***θ*** and ***ϕ*** matrices, which we then use to initialize/sample the strains of reads. In detail, once we condition upon ***θ*** and ***ϕ***, the random variables pertaining to different reads in our model become independent; hence we can sample each *z*_*m,n,r*_ independently using the observed *w*_*m,n,r*_ like so:

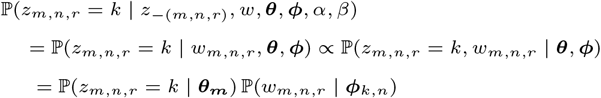

- **Hybrid SNP-LDA:** In this approach, the strain of a read with a mutation that uniquely identifies a known/reference strain is initialized to that strain, and all other reads are initialized to a random strain as described above. This reference-based approach (see Figure 1A.2) utilizes information about known strains during the initialization and iterative CGS phases of the algorithm to detect other minority strains more effectively; and is inspired by earlier topic-modeling works that used seed words capturing the underlying corpus to better detect rare topics [27, 28]. The premise here is that if information about most of the strains is known, then the proportion of the unknown minority strains can also be detected accurately.

We consider the barcoding SNP-alleles identified in [29] that discriminates nearly 91 *M. tb* strains as reference SNP-alleles. The reference SNP-alleles identified in this work are also incorporated into TBProfiler. Each of these reference SNP-alleles is a mutation that uniquely identifies the corresponding reference strain. The reference SNP-alleles present in each sample, the corresponding reference strains, and the hierarchical relationship between these strains are all utilized to assign strain identifiers to reads, both during initialization phase and during the iterative CGS inference phase (see details in Supplementary Section 1.2 and Algorithm 1 in Supplementary Information). Please note that hybrid SNP-LDA can also be run with other reference databases, which may include both unique and non-unique reference SNP-alleles (see Supplementary Section 1.2), but we assume a sufficient number of unique reference SNP alleles are present to reliably identify strains.

- **Summary of SNP-LDA variants, and hyperparameter choices:** We’ve seen three variants of SNP-LDA so far, and one more can arise from a combination of the two initialization heuristics. These four variants can be categorized into two reference-free and two reference-based (also known as hybrid) variants as follows: (i) SNP-LDA – a vanilla variant that initializes the strain of each read randomly; (ii) NMF SNP-LDA (also denoted (NMF, SNP-LDA)) – variant that initializes the strains of reads in the SNP-LDA model using the NMF outputs; (iii) (non-NMF) Hybrid SNP-LDA - a reference-based variant that initializes the strain of reads containing reference mutations/SNP-alleles to the corresponding reference strain and other reads randomly; and (iv) (NMF, hybrid SNP-LDA) – another reference-based variant that initializes the strain of reads containing mutations of a user-revealed reference strain *p* to *p* itself and other reads using the NMF outputs (this variant has only limited application in synthetic benchmarks where a reference strain in a sample is revealed, and the benchmark task is to find the remaining *de novo* strains in the sample and all strain proportions). Across the SNP-LDA model variants above, we employed 5 different hyperparameter combinations (1, 0.01), (0.01, *A*), (0.01, *B*), (*A*, 0.01) and (*B*, 0.01). The first element in each set corresponds to the *α* hyperparameter and the second element to the *β* hyperparameter (see Supplementary Section 1.5 for details of *A* and *B*). We used synthetic datasets LDAmix1 and LDAmix2 (see Section 2.5 for the description of datasets) to select the best model variant and hyperparameter configuration. We also considered a baseline approach, NMF-only, where NMF outputs are the final estimates of ***ϕ*** and ***θ***.

###### 2. Heuristic for choosing *K*

For a given input dataset of samples, the default Demixer (hybrid SNP-LDA) uses reference mutations in these samples to determine the total number of strains *K*. Specifically, *K* is set to the number of reference strains whose unique mutations (SNP-alleles) are present in the input samples (denoted *K*^*′*^), plus 2 additional strains to facilitate the detection of new strains. We use 2 as the default additional value under the assumption that the reference database is up-to-date and not too many new strains would have evolved considering the slower growth rate of TB bacteria [30]. The number of additional strains can be adjusted accordingly when applying Demixer to other microbial species. We use this *K* = *K*^*′*^ +2 heuristic for all our analyses, with the exception of model hyperparameter tuning where we have set *K* = 3 or *K* = 4 depending on the dataset characteristics. Note that hyperparameter tuning experiments involves some reference-free SNP-LDA variants, which do not have access to the reference database to calculate *K*^*′*^; hence we hard-coded the values of *K* for these experiments. In general, *K* for reference-free (non-hybrid) variants of SNP-LDA or NMF-only approach can be set by trying different values of *K* and choosing the one that performs best on an evaluation metric of interest.

###### 3. Weight heuristic

In a dataset with large number of samples, the number of reference SNP-alleles detected in a sample *m* (denoted *A*_*m*_) is likely to be a small fraction of the total number of SNP-alleles (called in all the samples, and estimated as *N* assuming that majority of the *N* SNPs are non-reference with single allele). To handle this imbalance, we give reference SNP-alleles a higher weight [31] based on the intuition that the aggregate sum of the read count of reference SNP-alleles should be similar to that of non-reference SNP-alleles. This weight (of at least 2 for reference SNP-alleles) is formulated as:

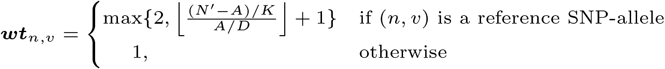

where 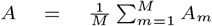 is the average number of reference SNP-alleles per sample, and *N* ^*′*^ = (Number of SNP-alleles)*/*2 ≈ *N* (as most SNPs are biallelic). The constant *D* is set by default to 3, which is derived empirically through the execution of our method on *in vitro* dataset across a range of constant values from 0 to 10, as it yielded a relative error that is comparable with the errors estimated using other values of D (see Supplementary Table S2). The weights determined using the above formula for the different datasets used in our study are shown in Supplementary Table S3. The CGS update formula in equation 2 on incorporating the above weights becomes:

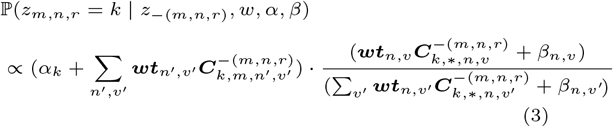

where *v* = *w*_*m,n,r*_.

###### 4. Parallelization heuristics

Since the running time of a CGS iteration can be quite high when the number of samples and reads are large, we tried parallelizing a CGS iteration. The “delayed count update” strategy used in an earlier WarpLDA model [32] can be used to parallelize CGS at the sample level. This delayed update CGS also enables assignment of strains to all reads of a SNP in a single step (using multinomial sampling based on the above equation 3, but using *C* instead of *C*^*−*(*m,n,r*)^), which is more efficient than sampling the strains of each read sequentially over multiple steps (using equation 3 as is). Our implementation of this parallel (delayed update) CGS and multinomial sampling heuristics is illustrated in Figure 2, and described in Algorithm 1 in Supplementary Information. These adaptations significantly improved the running time of one iteration of the CGS algorithm for processing all samples from *O*(*MNRK*) to *O* 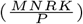, where *P* is the available number of processing cores (note *P* ≤ *M* as our parallelization is at the sample level). The space utilization of Demixer is *O*(*MNK*). The empirical memory usage and running time of Demixer for *in vitro* and real-world datasets are shown in Supplementary Table S4.

**Fig. 2.**
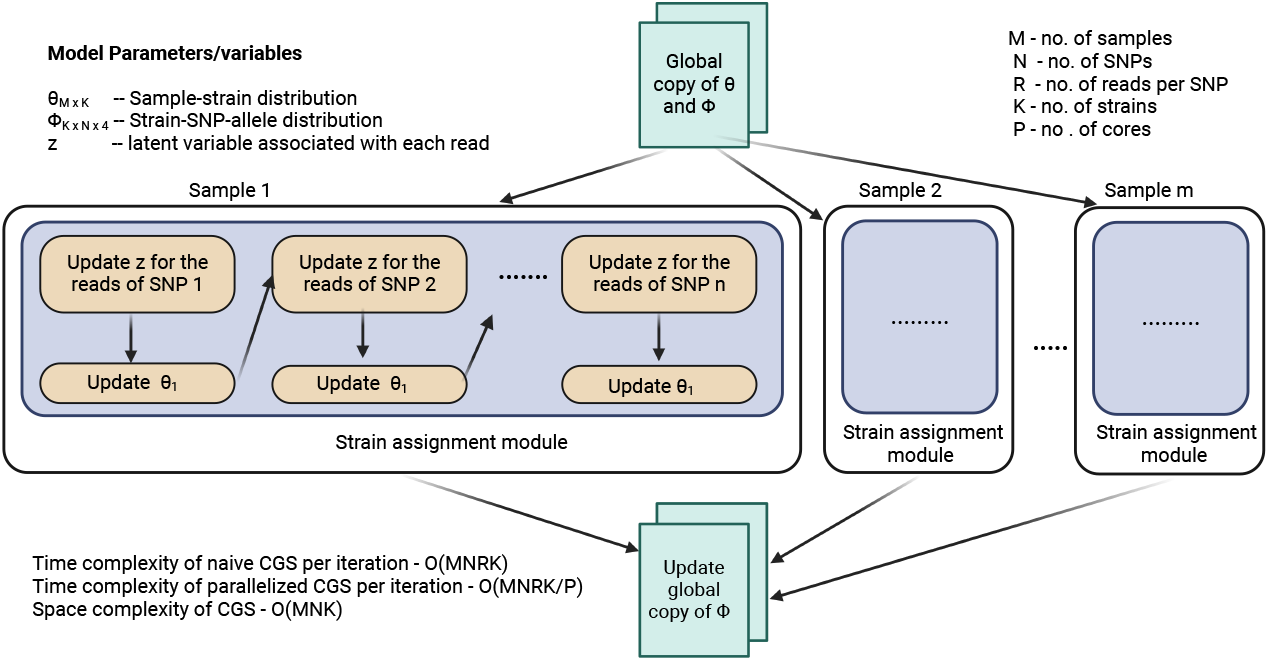
Heuristics for parallelizing the CGS algorithm. In one CGS iteration, all the samples are processed in parallel by maintaining local copies of the ***ϕ*** matrix. Each row in the ***θ*** matrix corresponds to a sample, and is updated after each SNP in that sample is processed, using the formula: ***θ***_*m,k*_ = ***C***_*k,m,∗,∗*_. Finally, the (global) ***ϕ*** matrix is updated after all the samples have been processed, using the formula: ***ϕ***_*k,n,v*_ = ***C***_*k,∗,n,v*_. Note that the ***θ*** and ***ϕ*** formula above correspond to unnormalized distributions, and can be normalized as shown in Algorithm 1 in Supplementary Information.

The different steps of SNP-LDA’s parameter estimation with initialization, choice of *K*, weight and parallelization heuristics are outlined in Algorithm 1 in Supplementary Information (also see Supplementary Section 1.7 for implementation details).

#### 2.3.3. Postprocessing/interpretation steps

To enhance the interpretability of our method’s predictions, we perform the following postprocessing steps on the strains inferred by Demixer:

- Mapping inferred to reference strains
- Fine-tuning the mapped strains using the lineage tree
- Quality checks using SNP plots/modes (*o*_*m*_ and *f*_*m*_ values) and *de novo* filtering
- Hierarchical clustering of the inferred strains, and *in vitro* dataset processing

Please refer Supplementary Section 1.3 for a detailed description of each postprocessing step, and Algorithm 2 in Supplementary Information for the order in which these steps are done. Supplementary Section 1.3 also provides the definition of the *f*_*m*_ value – briefly, *f*_*m*_ value of the *m*-th sample, expressed in terms of the number of SNPs, is used to decide if a mixed infection call is of low, medium or high confidence. This quality assessment is based on the distribution of minor allele proportions of the heterozygous SNPs in the sample.

### 2.4. Evaluation metrics

We use the following metrics to compare the performance of Demixer with other models and state-of-the-art methods on different benchmark datasets.

#### 1. Relative Error (RE)

We report relative error in simulated or *in vitro* datasets where we know the true/actual strain composition of a sample. The performance of a model/method on a given benchmark dataset is assessed by calculating the average relative error between the actual and predicted proportions for a strain *k* across all samples in that dataset (denoted *RE*_*k*_), and further averaging *RE*_*k*_ across all strains *k* ∈ {1, …, *K*} to obtain the final relative error (RE). Note that this averaging over all *K* strains is possible, since we assume that our mapping of inferred to ground-truth strains (following the same procedure explained above for mapping inferred to reference strains) follows a one-to-one relation. For all the datasets and methods where relative error is computed, this assumption held true. Note that in a given dataset, the relative error *RE*_*k*_ of the *k*^*th*^ strain can be computed as:

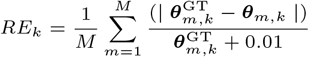

where 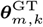 and ***θ***_*m,k*_ are respectively the actual (ground-truth) and predicted (estimated) proportion of strain *k* in sample *m*. The pseudocount 0.01 (minimum proportion of 1%) is added to the denominator to account for zero values of 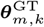.

#### 2. F1 score

QuantTB and Demixer are comparatively evaluated on benchmark datasets using the *F* 1 score outlined in [12] as the metric. The *F* 1 score of a sample quantifies how well a model/method can predict the actual strains present in the sample, which can then be averaged across all samples to get the *F* 1_*avg*_ score for a dataset. Before calculating the *F* 1 scores, all post-processing steps above excepting the last one related to hierarchical clustering are followed as described, but using the ground-truth strains of a benchmark dataset as the set of reference strains (along with its associated lineage tree). For each sample, the outcome of these post-processing steps is a one-to-one relation between the inferred and ground-truth strains; using which we can define a true positive (inferred strain mapped to a corresponding ground-truth strain), a false positive (unmapped inferred strain), and a false negative (unmapped ground-truth strain). These can in turn be used to calculate the F1 scores for the sample.

### 2.5. Datasets used in the analysis

Various synthetic and real-world datasets have been used in our study to tune and evaluate Demixer. Certain synthetic datasets were used for hyperparameter tuning alone (and not for evaluation). The remaining synthetic or *in vitro* benchmarking datasets generated using different procedures were used for evaluations. A brief note on these datasets follows (see Supplementary Section 1.4 for details and Supplementary Table S1 for dataset sizes).

- Datasets LDAmix1 and LDAmix2 are synthetic datasets created using our SNP-LDA model’s generative process to evaluate Demixer’s potential in delineating the mixed strains under different hyperparameter combinations and deciding if a simple hyperparameter value would work well in further analyses.
- To mimic the benchmark dataset used to evaluate QuantTB, we generated two standard datasets (ART-TBmix1 and ART-TBmix2) as per the simulation procedure described in [14][12] using ART simulator [33]. Both ART-TBmix1 and ART-TBmix2 consists of 800 samples at four distinct levels of coverage: 10x, 20x, 90x-10x, and 70x-30x, each with 200 samples. A sample in 10x and 20x datasets is mixed with four strains, whereas two strains are mixed in 90x-10x and 70x-30x datasets using the corresponding coverage levels. The strains in ART-TBmix1 samples are chosen from 2,165 strains in QuantTB’s [12] reference database, and that in ART-TBmix2 are chosen from 89 strains from Robust Barcoding database [29] (ignoring lineage 4 and 4.9 strains).
- Another dataset ARTmix was generated to evaluate the performance of Demixer in detecting *de novo* strains, when information about the other strains are revealed in the reference database. Each of the 50 samples in this dataset comprises either 1, 2, or 3 strains, which are selected from a set of 7 strains obtained by replacing 100 SNPs in the reference H37Rv genome.
- An *in vitro* dataset of 48 samples obtained by experimentally mixing the DNA of two *M. tb* samples (belonging to lineages 1-4) in one of these proportions: 0.7/0.3, 0.90/0.10, 0.95/0.05, and 1.00/0.00. This realistic benchmark dataset from an earlier study [13] is used to compare Demixer with other published methods. We observed a few discrepancies on examining the ground-truth information pertaining to *in vitro* dataset and have made changes to the lineages of the strains present in 8 mixed samples. Specifically, CAS1-Delhi is assigned its original lineage 3 [34], and LAM11-ZWE is assigned lineage 4 [35]. The modified lineages and the lineages identified by Demixer and QuantTB are reported in Supplementary File D1.
- Two real-world datasets, comprising 1,963 TB isolates obtained from patients in Malawi [13] and 12287 TB isolates obtained across 23 countries by CRyPTIC [36] has been used to detect mixed infections. The CRyPTIC dataset is also examined to assess the impact of mixed infection on drug resistance.

## 3. Results

### 3.1. Model selection and hyperparameter tuning using synthetic datasets

The selection of appropriate hyperparameters for a probabilistic model such as LDA can play a role in improving its performance in text mining tasks [37]. Since our SNP-LDA is a newly proposed LDA model, we wanted to first check if the model is sensitive or robust to the choice of hyperparameters and other settings. We considered different variants of SNP-LDA model resulting from different hyperparameter settings (see Supplementary Section 1.5) and initialization heuristics (see Methods), and compared their performances on two synthetic datasets (see Figure 3).

**Fig. 3.**
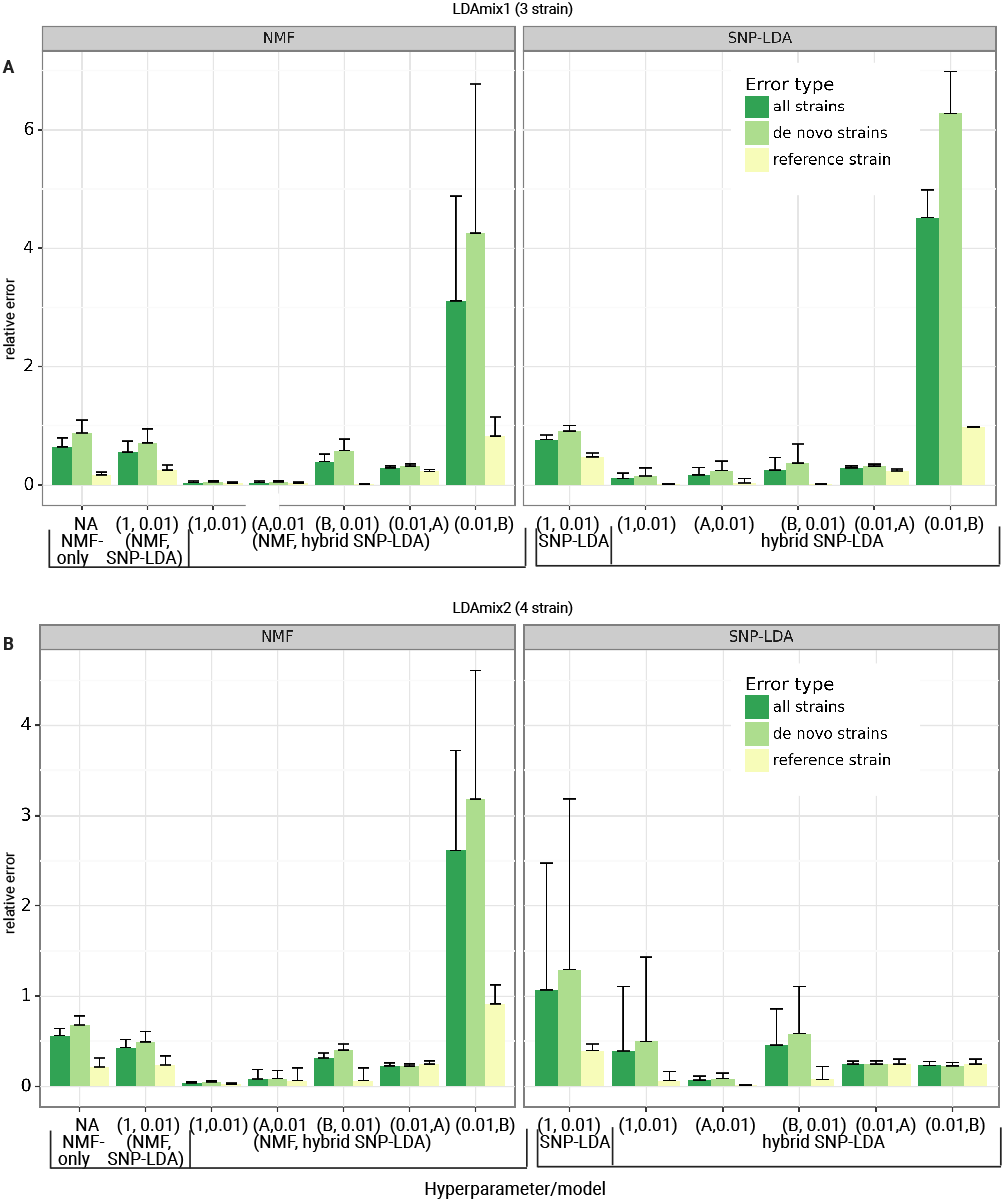
Performance of different SNP-LDA model variants on two synthetic datasets: Model variants are indicated in a pairwise tuple format (see Methods, Section 2.3.2). A) Performance on the three-strain mixture dataset LDAmix1. B) Performance on the four-strain mixture dataset LDAmix2. Each dataset has been generated 10 times (i.e., in 10 runs using different random seeds). The x-axis indicates the different methods tested (with their hyperparameter/model combinations; see Supplementary Section 1.5), and the y-axis indicates the relative error between the actual and predicted strain proportions averaged across the 10 runs (with error bar overall length being the standard deviation across the 10 runs). The error type indicates the error in determining all the strains, solely reference strains, and only the *de novo* strains in the samples within each dataset.

Using relative error as the evaluation measure, we observed that all model variants, except the one with hyperparameter (0.01, *B*) (see Methods), were able to accurately predict the reference strain proportions. For predicting *de novo* strain proportions, we found the error to be lower for most hybrid SNP-LDA combinations, indicating the effectiveness of using information (mutations) of known strains on top of the SNP-LDA model. Also, the symmetric hyperparameter setting (1, 0.01) was similar or better in performance compared to the other asymmetric hyperparameter settings; so we will use the former for all future analyses.

The (NMF, hybrid SNP-LDA) combined models leaving out the one with hyperparameter (0.01, B) have lower relative error than NMF-only models (indicating the benefit from probabilistic treatment of uncertainty in SNP-LDA) and non-NMF models as well. But, (NMF, hybrid SNP-LDA) models have limited applicability beyond synthetic benchmarks as discussed in Methods; further it can be challenging to assign reference SNP-allele labels to NMF-inferred strains from real-world data. Therefore, we use the (non-NMF) hybrid SNP-LDA model with hyperparameter (1, 0.01) as the default model for Demixer for all subsequent analyses. We have also compared our Demixer to the traditional LDA model on a 2-strain dataset when no reference strains are known (see Supplementary Section 1.6), and the better performance of our model on this dataset (Supplementary Figure S2) supports why a new model such as ours is needed for estimating strain proportions in WGS samples.

### 3.2. Demixer performs better than QuantTB in the *in vitro* dataset

We compared the performance of our Demixer with state-of-the-art methods QuantTB and SplitStrains using the *in vitro* dataset consisting of 48 samples obtained by experimentally mixing two pure samples in different proportions. In this work, QuantTB and SplitStrains are run on the samples using their corresponding default settings and parameter values. We use the hierarchical clustering tree of the strains inferred by Demixer (Figure 4A) to group the strains into two clusters (see Supplementary Section 1.3), and thereby estimate the major and minor strain proportions. These predicted proportions by Demixer are very similar to the actual proportions and the associated relative error (RE) is much lower than QuantTB and comparable to SplitStrains (Figure 4B). Demixer was also better than QuantTB in inferring the lineages of strains correctly (Figure 4C; see also Supplementary File D1 for the inferred lineages). Demixer inferred strains related to lineages 1–4 correctly in most of the samples, except for one unmapped strain in a sample (that is closer to lineage 2 in the hierarchical tree, and explained in detail in Supplementary File D1).

**Fig. 4.**
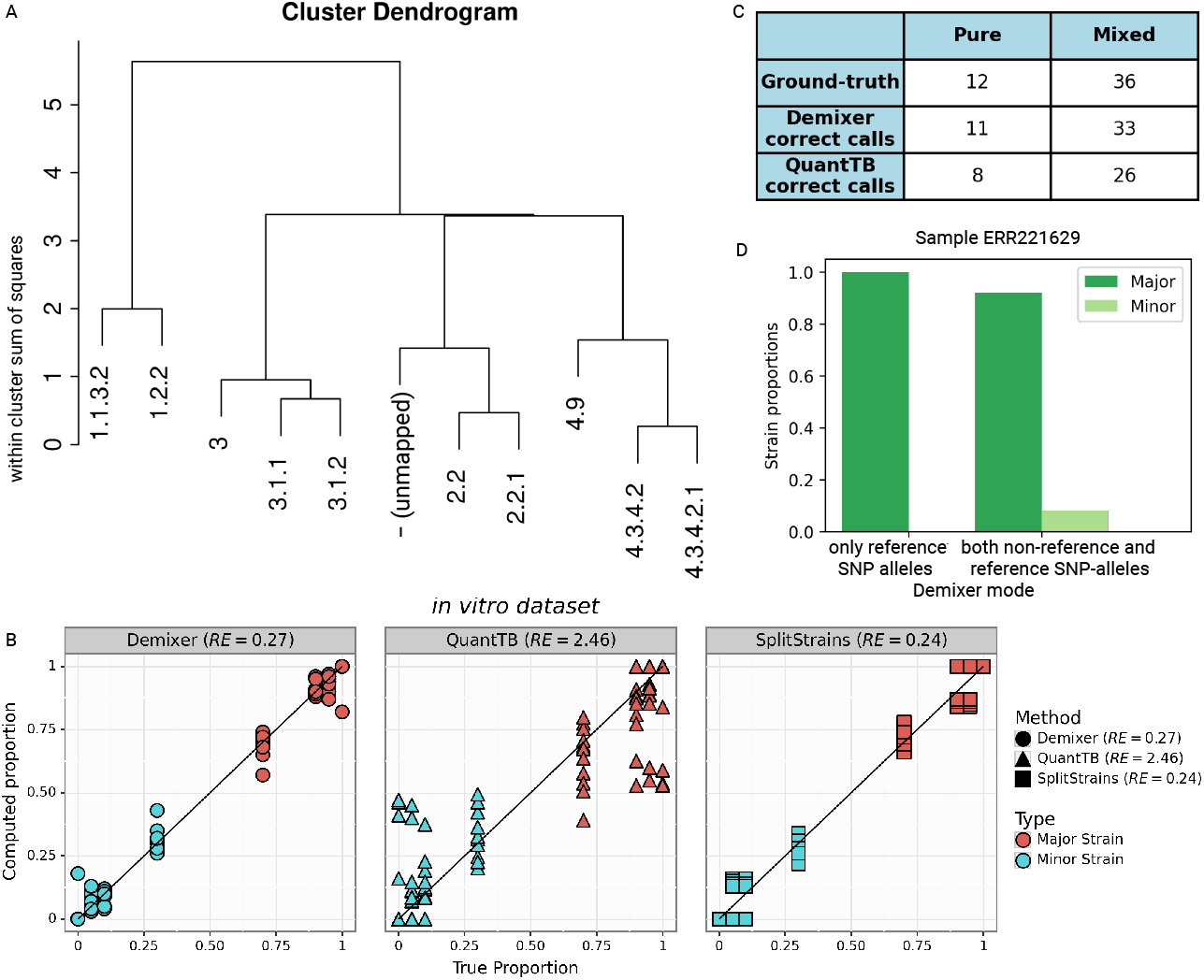
Demixer vs. other methods on *in vitro* dataset: A) represents the hierarchical relationship between the strains inferred by Demixer in the *in vitro* dataset. B) compares the predicted vs. actual proportions of the major/minor strain in each sample. C) represents the number of samples for which the strains inferred by Demixer (or QuantTB) mapped correctly to the lineages present in the sample. D) represents the strain proportions identified by Demixer for the sample ERR221629, which is actually a mixed sample with ground-truth strain proportions 0.90/0.10. SplitStrains can detect only proportions and not strain identities, so it is not shown in panel C.

Demixer uses both reference and non-reference SNP-alleles to perform its inference, and this offers an advantage as demonstrated by Demixer correctly identifying the sample ERR221629 (with ground truth proportion 0.90/0.10) as mixed; because when only reference SNP-alleles were used, Demixer incorrectly classified this sample as pure (Figure 4D).

### 3.3. Evaluating Demixer on QuantTB and novel-strain-identification benchmarks

To enable further comparison of a multi-sample analysis method like Demixer with a per-sample-analysis tool like QuantTB, we evaluated them on the same (or similar) benchmarks published in the QuantTB work. The benchmark datasets named ART-TBmix1,2 in our study are obtained by artificially mixing simulated samples at different coverages 10x, 20x, 90x-10x, and 70x-30x (see Supplementary Section 1.4). The *F* 1_*avg*_ score of Demixer is comparable to that of QuantTB for the ART-TBmix1 dataset, whereas better than QuantTB for the ART-TBmix2 dataset (Figure 5B). Note that the *F* 1_*avg*_ scores of Demixer are greater than 0.90 at all coverage levels of ART-TBmix{1,2}. Demixer has relatively lower performance at 10x coverage level in ART-TBmix1, which may be due to the low coverage data not being sufficient to learn the parameters of the probabilistic model underlying Demixer.

**Fig. 5.**
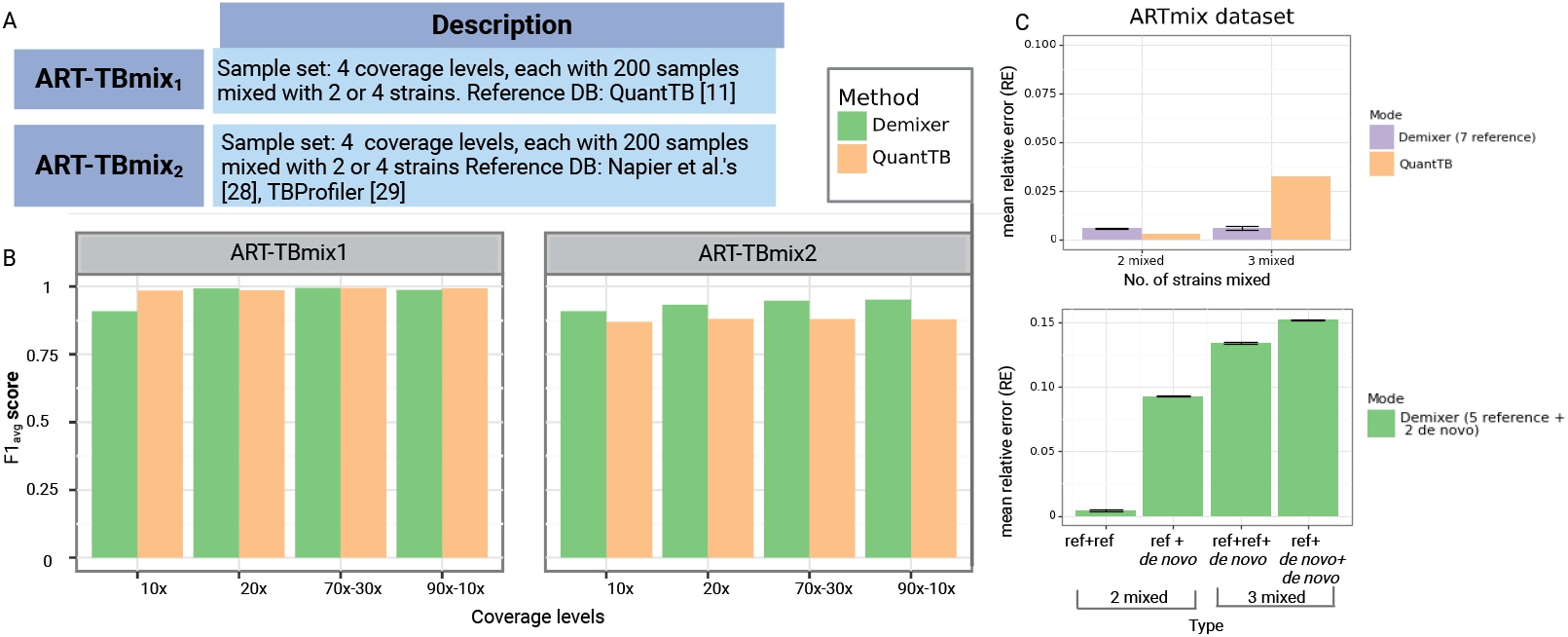
Comparison of Demixer vs. QuantTB: A) The table highlights the number of samples present in the datasets ART-TBmix1 and ART-TBmix2. The strains of ART-TBmix1 and ART-TBmix2 are generated using the SNP-alleles of strains based on QuantTB [12] and Napier et al.’s [29] works respectively. B) *F* 1_*avg*_ scores obtained by running Demixer and QuantTB on ART-TBmix1 and ART-TBmix2 datasets. These scores along with the related precision and recall measures are in Supplementary Tables S5 and S6. C) Performance of the two modes of Demixer on ARTmix data. The y-axis of both the top and bottom plots represents the mean RE (see Methods), with error bar overall length being twice the standard deviation across the 10 runs of Demixer (each involving a different random seed). The x-axis of the top plot corresponds to the number of mixed strains present in ARTmix samples and the x-axis of bottom plot correspond to the different combinations of reference and *de novo* strains in the ARTmix samples.

We also compared Demixer and QuantTB using the ARTmix dataset consisting of 18 pure and 32 mixed samples (see Supplementary Section 1.4) in “all references known” mode. In this mode, the SNP-alleles of the strains used to generate the ARTmix samples were revealed in the form of a reference database to both Demixer and QuantTB (see Methods and Supplementary Section 1.4). As ARTmix includes mixed samples with both 2 and 3 strains, we have computed the mean RE for each case by including only the corresponding samples. Though Demixer’s mean RE is slightly higher than that of QuantTB in samples mixed with 2 strains, they are still comparable as the relative errors are very low (top plot of Figure 5C).

While QuantTB is a state-of-the-art method for estimating TB strain proportions, it is fully reference-based and cannot identify novel strains. Our Demixer however can identify new strains due to its hybrid (reference-plus-*de novo*) design. To evaluate how well Demixer can identify new strains, we ran it in “few references known” (*de novo*) mode on the ARTmix dataset. In this mode, the SNP-alleles of only five strains are revealed to Demixer. Demixer’s RE in *de novo* mode (bottom plot of Figure 5C) is, as expected, higher than Demixer in “all references known” mode and QuantTB. However, the RE is at most 0.15, and so Demixer in *de novo* mode can identify strains whose SNP-alleles are not in the reference database with reasonable accuracy.

### 3.4. Demixer detects new mixed samples in Malawi dataset

In the previous section, we have seen how Demixer can detect *de novo* strains within a synthetic dataset. To assess the same in a real-world setting, we consider the Malawi samples comprising major lineages (1–4) of the species *M. tb* and another TB-related species *M. bovis*. We specifically applied Demixer on these samples after hiding the unique SNP-alleles of *M. bovis* strain in the reference database. The SNP-alleles corresponding to 40 strains are found in the sample set, so the model parameter *K* is set to 42 (see Methods). TBProfiler identifies 5 pure *M. bovis* samples and one sample mixed with *M. bovis* and lineage 3 strain in the Malawi dataset. The proportions determined by Demixer for those 6 samples containing the unmapped strain T41 (Figure 6A - B) indicate that Demixer can delineate the proportions even after the exclusion of *M. bovis* SNP-alleles from the reference database. The new strain T41 was detected only in the above six *M. bovis* samples, and the strain’s proportion in the five pure *M. bovis* samples were also correctly identified as 100%. The 62 mutations of the T41 strain, as inferred from the Strain-SNP-allele distribution provided by Demixer is in agreement with the reference mutations of *M. bovis* strain. These results indicate that Demixer can infer the mutational profile as well as the proportion of the *de novo* strain (*M. bovis* in this case) accurately.

**Fig. 6.**
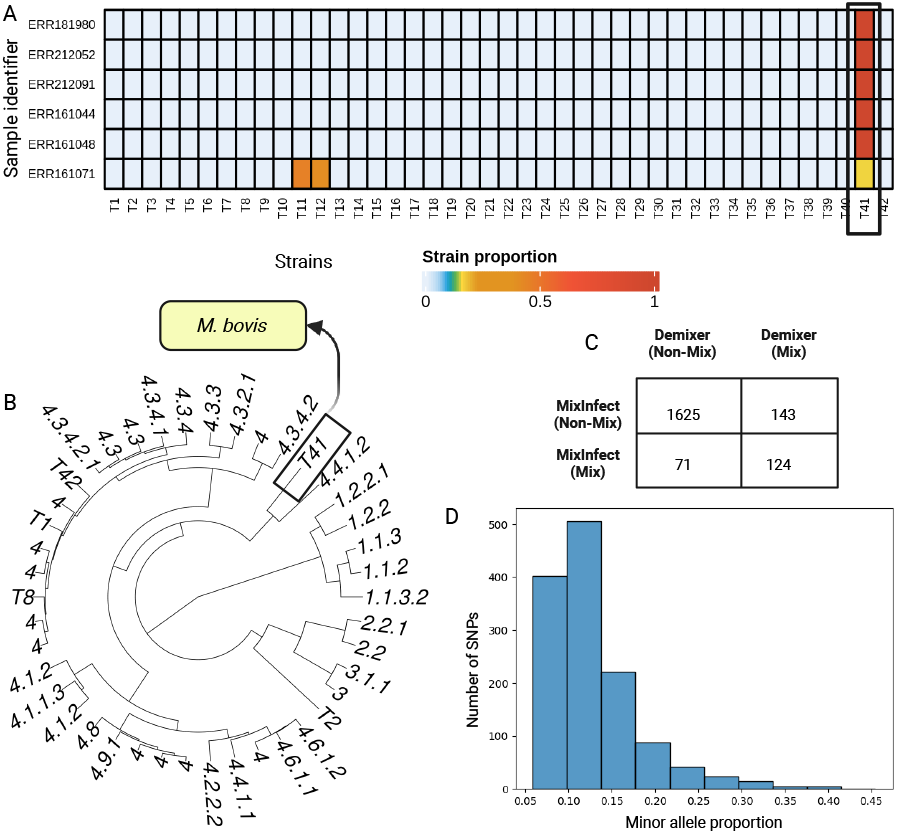
Demixer results on Malawi dataset: A) represents the Demixer-estimated strain proportions of samples containing *M. bovis* strains. Here, Demixer is run by hiding the SNP-alleles relevant to *M. bovis* strain. B) represents the hierarchical clustering of the inferred strains. The strain number T41 corresponds to the new strain identified by Demixer, which is distant from the lineages 1, 2, and 3. C) shows the number of samples identified as mixed and non-mixed by Demixer and MixInfect. D) represents the SNP plot of sample ERR161149 identified as mixed by Demixer and non-mixed by MixInfect. The x-axis refers to the proportion of the minor alleles in a heterozygous SNP and y-axis refers to the frequency of such SNPs with that minor allele proportion.

Next, we applied Demixer to the 1,963 Malawi samples after including all the strains in the reference database. Demixer was able to identify known (124) as well as novel (143) mixed infection samples, relative to previously reported mixed samples [13] (see Figure 6C and Supplementary File D2). The previously reported calls were made by MixInfect, a tool similar to SplitStrains in terms of estimating the proportion but not the mutational profile of strains. To higlight an example of a novel mixed infection sample, consider ERR161149 - Demixer identifies this sample as a mixture of strains 4.3.4.2.1 (95%) and 4.4.1.2 (5%), whereas MixInfect calls this sample as pure. When checking this sample using a SNP plot (Figure 6D), we found clear evidence for a mixture of strains (due to the presence of 1309 heterozygous SNPs in the sample with mode *o*_*m*_ being equal to 11%). When checking all 143 novel mixed calls using a SNP plot related confidence measure *f*_*m*_ (see Methods), we could classify all but 5 of them as medium/highly confident mixed calls (Supplementary Figure S3A); further, their *f*_*m*_ values taken together was higher than the *f*_*m*_ values of the 71 samples called as mixed by MixInfect but not Demixer (Wilcoxon ranksum test one-sided p-value 2.21 *×* 10^*−*25^, with associated statistic value of 9489; see also Supplementary Figure S3B). Taken together, our multi-sample analysis method Demixer can detect mixed infection with high sensitivity in samples obtained from a clinical setting.

### 3.5. Demixer infers global frequency of mixed infections and its association to drug resistance

We applied Demixer to CRyPTIC, a large dataset of *M. tb* genomes of isolates collected from patients distributed across 23 countries [38], in order to estimate the frequency of mixed infection across the globe. To detect mixed infection confidently in a clinical setting and to assess its relation to drug resistance, we focus only on Demixer’s high-confidence calls on the CRyPTIC dataset (as determined using the *f*_*m*_ and *o*_*m*_ measures; see Methods and Supplementary Figure S4A). Demixer identified 385 (3.3% of the total 12287) isolates to be mixed infection with high confidence (see Figure 7A). When testing overlap against the 6813 CRyPTIC isolates resistant to at least one TB drug, we found that 225 of these 385 mixed infection isolates are drug-resistant. This overlap is marginally significant (hypergeometric p-value 0.1253). However as shown in Figure 7A, the statistical association between mixed infection and drug resistance becomes stronger as the stringency for high-confidence mixed infection calls by Demixer are increased progressively (from 75^th^ to 95^th^ percentiles of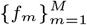, with hypergeometric p-value 0.0099 for the most stringent threshold).

**Fig. 7.**
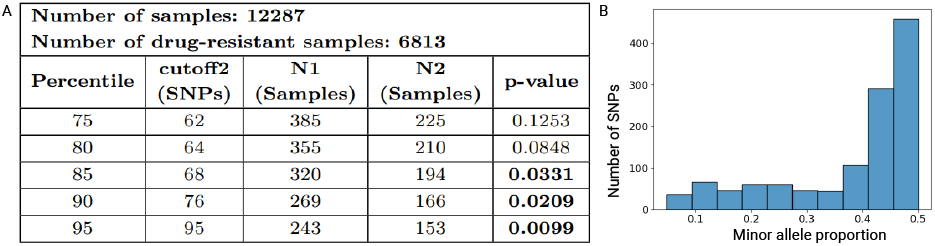
Association between mixed infection and drug-resistance in CRyPTIC samples. A) Hypergeometric test p-values to assess the association between drug resistance and mixed infection are shown, using high-confidence mixed infection calls based on different *f*_*m*_ and *o*_*m*_ thresholds. *N* 1 refers to the number of mixed samples whose *f*_*m*_ *>* cutoff2 and *o*_*m*_ *>* 0.05, and *N* 2 refers to a subset of the samples accounted for in *N* 1 that are also resistant to at least one drug. B) SNP plot of the mixed infection sample resistant to all the 13 tuberculosis drugs. The interpretation of SNP plot is same as in Figure 6D. See Supplementary File D3 for the estimated lineages and proportions of all CRyPTIC samples.

In addition to revealing global patterns of mixed infection and its link to drug resistance in the CRyPTIC dataset, Demixer could also reveal useful information about each individual sample. For instance, of the 2 CRyPTIC samples that are resistant to all 13 drugs, Demixer identified one of them as a pure sample of lineage 2.2.1 and the other as a mixture of strains belonging to lineages 2.2.1 (44%) and 4.2.1 (56%). The SNP plot of the latter mixed sample is shown in Figure 7B. As another application of Demixer, we also collated its predictions to calculate the percentage of mixed infection samples that are resistant separately to each antituberculous drug (see Supplementary Figure S4B). Finally, we analyzed a small subset of samples from the Malawi dataset using the ***ϕ*** obtained by running Demixer on the CRyPTIC dataset, emphasizing the efficacy of the method for real-time analysis (see Supplementary File D4 for the inferred strains and proportions on the Malawi subset). These applications together show that Demixer is a useful tool for predicting mixed infection and assessing its link to drug-resistance.

## 4. Discussion

This work presents Demixer, a new probabilistic approach based on topic modeling to aid in the detection of mixed strains present in microbial samples. Such modeling has enabled a multi-sample analysis, which is first of its kind to estimate identity and proportion of mixed strains from WGS data, with comparable or better performance over other approaches. The utilization of information about known strains gives Demixer the advantage of reference-based approaches, whereas the probabilistic modeling along with multi-sample analysis facilitate the identification of *de novo* strains. The parallelization heuristic allows Demixer to scale to large datasets (containing 1500 or more samples). Demixer has been extensively evaluated on different synthetic benchmarks and real-world datasets. For instance, Demixer employing simple hyperparameters (chosen from a systematic tuning approach using synthetic LDAmix* datasets) is effective in estimating strain proportions on the *in vitro* benchmark and other synthetic datasets. Demixer’s results on the Malawi and CRyPTIC dataset demonstrate the benefits of using prior information in determining the identity of strains in samples obtained from a clinical setup. They also point to a scenario where Demixer parameters learnt from a large dataset like CRyPTIC can be used to infer strains in new samples in a smaller dataset refer Supplementary Section 1.8. Demixer is generalizable and can be applied to other bacterial or microbial species as well.

The strain estimations by our Demixer and the previously published MixInfect methods are agreeable in 89% of the samples, and the disagreements could be attributed to differences in the variant calling or preprocessing workflows, and the mixed infection detection methodology (e.g., single-vs. multi-sample analysis). The detection of 3.1% mixed infections (after post-processing using default cutoffs) in the CRyPTIC dataset is not too far from the 1.5% mixed infections reported in [38]. Additionally, Demixer identified four *M. orygis* samples, which were not reported in [38]. When comparing Demixer with other methods on real-world datasets, we resorted to SNP plots and associated measures (*f*_*m*_ values) to validate the strains predicted by a method. We used these proxy validation measures to compare methods, since ground-truth experimentally validated mixed-infection datasets (like the *in vitro* dataset) are scarce. Once more such datasets become available, we can better address the uncertainty around whether the presence of SNPs from two different strains in a sample truly indicates mixed infection, and if so, what is the minimum number of such SNPs required to be present.

Huang et al. [39] emphasize the importance of identifying the lineages of strains in an individual before initiating the treatment, as mixed infections due to nontuberculous mycobacteria (NTM) and *M. tb* can impact treatment outcomes. Therefore, a reference database consisting only of *M. tb* lineages may not be sufficient to identify mixed infections due to *M. tb* and NTM. The increasing trend of TB caused by *M. bovis*, with its natural resistance to pyrazinamide, further complicates matters [40]. Demixer has the potential to identify such NTM strains even if they are not present in the reference database, as they are dissimilar from *M. tb* sequences. Our results on synthetic and real-world datasets demonstrated the effectiveness of Demixer in disentangling dissimilar *de novo* strains. The importance of developing new techniques to identify novel strains has also been highlighted by the authors of the recently published Fastlin tool [41].

Regarding limitations of Demixer, one potential concern with our approach is that a model that is tuned on one dataset (LDAmix* in our case) may not work well on other datasets; however this was not the case in the applications of Demixer we have tried till now. Another concern could be about the completeness of the reference database that we use with Demixer, even though we’ve taken care to choose a widely-used reference database. For instance, the latest subtypes of lineage 2 are still missing in the reference database, and this could mislead inferences of Demixer on samples containing these subtypes. Demixer could also be misled about closely related strains that have insufficient number of SNP-alleles separating them in the reference database. This underscore the need for efforts to continuously update the reference database used with Demixer. A final concern is that, like other LDA models, Demixer requires careful selection of the number of strains *K* to achieve meaningful and interpretable results. A high *K* value may mislead the model to declare a sub-pattern present in a lineage as a new strain, thereby leading to unnecessary fragmentation of strains. We carefully choose *K* using the *K* = *K*^*′*^ + 2 heuristic, where *K*^*′*^ captures only the reference strains whose SNP-alleles are in the input samples and the extra 2 enables detection of *de novo* strains. Through this heuristic and via post-processing (hierarchical clustering) of sub-strains into fewer strains, we mitigated the fragmentation issue in complex real-world TB datasets. Future work could focus on improving Demixer by developing better heuristics for selecting *K* and additionally also for assigning weights to reference SNP-alleles (since Demixer performance can be sensitive in certain datasets to the current weighting heuristics; see Supplementary Figure S5).

Demixer represents a novel systematic approach for delineating the mixed strains in microbial samples, as demonstrated here in its applications to diverse datasets. These results encourage different future extensions of Demixer. One interesting future extension could be to model the hierarchical relationship between the inferred strains directly into the underlying SNP-LDA model of Demixer and see if the performance is further improved. In the current implementation, Demixer incorporates an input lineage tree (and the assumption that SNP-alleles present in a parent strain are also present in descendant strains) to infer hierarchically related strains present in a sample. Another fruitful extension of Demixer could be to analyze metagenomic reads for strain-level identification [42]. Demixer in its current and extended forms holds promise to reveal new insights into mixed infection diagnosis and treatment.

## 5. Acknowledgments

We thank members of our BIRDS (Bioinformatics and Integrative Data Science) and IBSE (IBSE Centre for Integrative Biology and Systems medicinE) (IBSE) research groups for their valuable inputs during the presentations of this work. We thank Dr. S. Sivakumar from National Institute for Research in Tuberculosis, Chennai (NIRT) for his valuable suggestions on interpreting the model outputs. We also extend our thanks to Dr. Ranjeeta Menon from Monash University for her insightful feedback on the manuscript. The tools from biorender.com were used to create Figures 1, 2 and S1 and to edit other figures in the manuscript. The research presented in this work was supported by Wellcome Trust/DBT grant IA/I/17/2/503323 awarded to MN.

