## Supplemental Information for "Demixer: A probabilistic generative model to delineate different strains of a microbial species in a mixed infection sample"


1 Supplementary Methods **1**
- 1.1 CGS inference and parameter estimation ... 1
- 1.2 From a reference database to global/sample-specific dictionaries ... 2
- 1.3 Postprocessing/interpretation steps ... 3
- 1.4 Datasets ... 4
- 1.5 Model selection and hyperparameter tuning ... 5
- 1.6 Comparison of Demixer and traditional LDA ... 5
- 1.7 Implementation details ... 5
- 1.8 Using Demixer for analysing new sample(s) ... 5

2 Supplementary Algorithms **7**

3 Supplementary Tables **9**

4 Supplementary Figures **11**

5 Supplementary Files **13**


### 1. Supplementary Methods

#### 1.1. CGS inference and parameter estimation

The probability of assigning a strain to a read (i.e., sampling $z_{m,n,r}$) given the strain assignment of all other reads (denoted by $z_{-(m,n,r)}$) is derived [2] as below:

$$\begin{aligned}\mathbb{P}(z_{m,n,r} = k \mid z_{-(m,n,r)}, w, \alpha, \beta) &\propto \mathbb{P}(z_{m,n,r} = k, z_{-(m,n,r)}, w \mid \alpha, \beta) \\ &= \mathbb{P}(z, w \mid \alpha, \beta) \end{aligned} \tag{1}$$

Now, we derive $\mathbb{P}(z, w \mid \alpha, \beta)$ using our model structure (i.e., conditional independence assumptions implied by the model) as follows:

$$\begin{aligned}\mathbb{P}(z, w \mid \alpha, \beta) &= \mathbb{P}(z \mid \alpha) \times \mathbb{P}(w \mid z, \beta) \\ &= \int \mathbb{P}(\boldsymbol{\theta} \mid \alpha)\ \mathbb{P}(z \mid \boldsymbol{\theta})\ \mathrm{d}\boldsymbol{\theta} \times \int\ \mathbb{P}(\boldsymbol{\phi} \mid \beta)\ \mathbb{P}(w \mid \boldsymbol{\phi}, z)\ \mathrm{d}\boldsymbol{\phi} \\ &= \int \prod_{m=1}^{M} \mathbb{P}(\boldsymbol{\theta}_m \mid \alpha) \prod_{n=1}^{N}\prod_{r=1}^{R} \mathbb{P}(z_{m,n,r} \mid \boldsymbol{\theta}_m)\ \mathrm{d}\boldsymbol{\theta} \times \int \prod_{n=1}^{N}\left(\prod_{k=1}^{K} \mathbb{P}(\boldsymbol{\phi}_{k,n} \mid \beta_n)\right) \prod_{m=1}^{M}\prod_{r=1}^{R} \mathbb{P}(w_{m,n,r} \mid \boldsymbol{\phi}_{z_{m,n,r},n})\ \mathrm{d}\boldsymbol{\phi} \\ &= \left(\prod_{m=1}^{M}\int \mathbb{P}(\boldsymbol{\theta}_m \mid \alpha) \prod_{n=1}^{N}\prod_{r=1}^{R} \mathbb{P}(z_{m,n,r} \mid \boldsymbol{\theta}_m)\ \mathrm{d}\boldsymbol{\theta}_m\right) \times \left(\prod_{n=1}^{N}\int \left(\prod_{k=1}^{K} \mathbb{P}(\boldsymbol{\phi}_{k,n} \mid \beta_n)\right) \prod_{m=1}^{M}\prod_{r=1}^{R} \mathbb{P}(w_{m,n,r} \mid \boldsymbol{\phi}_{z_{m,n,r},n})\ \mathrm{d}\boldsymbol{\phi}_n\right)\end{aligned} \tag{2}$$

Letting $V = \{A, C, G, T\}$, expanding the discrete distribution $\mathbb{P}(w \mid \boldsymbol{\phi}, z)$ using the indicator function $\mathbb{1}$ (defined in the main text), and grouping the resulting terms by $q \in V$ and by the strains $k$ as in [2], we can simplify some of the terms in the above equation as follows:

$$\begin{aligned}\prod_{m=1}^{M}\prod_{r=1}^{R} \mathbb{P}(w_{m,n,r} \mid \boldsymbol{\phi}_{z_{m,n,r},n}) &= \prod_{m=1}^{M}\prod_{r=1}^{R}\prod_{q\in V} \boldsymbol{\phi}_{z_{m,n,r},n,q}^{\mathbb{1}(w_{m,n,r}==q)} \\ &= \prod_{m=1}^{M}\prod_{r=1}^{R}\prod_{q\in V}\prod_{k=1}^{K} \boldsymbol{\phi}_{k,n,q}^{\mathbb{1}(w_{m,n,r}==q \text{ AND } z_{m,n,r}==k)} \\ &= \prod_{q\in V}\prod_{k=1}^{K} \boldsymbol{\phi}_{k,n,q}^{\sum_{m=1}^{M}\sum_{r=1}^{R} \mathbb{1}(w_{m,n,r}==q \text{ AND } z_{m,n,r}==k)} \\ &= \prod_{k=1}^{K}\prod_{q\in V} \boldsymbol{\phi}_{k,n,q}^{\boldsymbol{C}_{k,*,n,q}}\end{aligned} \tag{3}$$

Similarly,

$$\prod_{n=1}^{n}\prod_{r=1}^{R}\mathbb{P}(z_{m,n,r}\mid\boldsymbol{\theta}_m)=\prod_{k=1}^{K}\boldsymbol{\theta}_{m,k}^{\boldsymbol{C}_{k,m,*,*}} \tag{4}$$

Substituting Equations 3, 4 into Equation 2, we get:

$$\begin{aligned}\mathbb{P}(z,w\mid\alpha,\beta)&=\left(\prod_{m=1}^{M}\int\mathbb{P}(\boldsymbol{\theta}_m\mid\alpha)\prod_{k=1}^{K}\boldsymbol{\theta}_{m,k}^{\boldsymbol{C}_{k,m,*,*}}\,\mathrm{d}\boldsymbol{\theta}_m\right)\times\left(\prod_{n=1}^{N}\int\left(\prod_{k=1}^{K}\mathbb{P}(\boldsymbol{\phi}_{k,n}\mid\beta_n)\right)\prod_{k=1}^{K}\prod_{q\in V}\boldsymbol{\phi}_{k,n,q}^{\boldsymbol{C}_{k,*,n,q}}\,\mathrm{d}\boldsymbol{\phi}_n\right)\\&=\left(\prod_{m=1}^{M}\int\mathbb{P}(\boldsymbol{\theta}_m\mid\alpha)\prod_{k=1}^{K}\boldsymbol{\theta}_{m,k}^{\boldsymbol{C}_{k,m,*,*}}\,\mathrm{d}\boldsymbol{\theta}_m\right)\times\left(\prod_{n=1}^{N}\prod_{k=1}^{K}\int\mathbb{P}(\boldsymbol{\phi}_{k,n}\mid\beta_n)\prod_{q\in V}\boldsymbol{\phi}_{k,n,q}^{\boldsymbol{C}_{k,*,n,q}}\,\mathrm{d}\boldsymbol{\phi}_{k,n}\right)\\&=\left(\prod_{m=1}^{M}\int\frac{1}{B(\alpha)}\prod_{k=1}^{K}\boldsymbol{\theta}_{m,k}^{\alpha_k-1+\boldsymbol{C}_{k,m,*,*}}\,\mathrm{d}\boldsymbol{\theta}_m\right)\times\left(\prod_{n=1}^{N}\prod_{k=1}^{K}\int\frac{1}{B(\beta_n)}\prod_{q\in V}\boldsymbol{\phi}_{k,n,q}^{\beta_{n,q}-1+\boldsymbol{C}_{k,*,n,q}}\,\mathrm{d}\boldsymbol{\phi}_{k,n}\right)\\&=\left(\prod_{m=1}^{M}\frac{B\left(\{\alpha_k+\boldsymbol{C}_{k,m,*,*}\}_{k=1}^{K}\right)}{B\left(\{\alpha_k\}_{k=1}^{K}\right)}\right)\times\left(\prod_{n=1}^{N}\prod_{k=1}^{K}\frac{B\left(\{\beta_{n,q}+\boldsymbol{C}_{k,*,n,q}\}_{q\in V}\right)}{B\left(\{\beta_{n,q}\}_{q\in V}\right)}\right)\end{aligned}$$

In the last two steps above, we had expanded the Dirichlet distributions $\mathbb{P}(\boldsymbol{\theta}_m\mid\alpha)$ and $\mathbb{P}(\boldsymbol{\phi}_{k,n}\mid\beta_n)$ and used the normalizing constants of the Dirichlet distribution (the multivariate beta function $B(.)$). Substituting above expression into Equation 1, and simplifying further by expanding the beta functions and dropping constant terms as in [2], yields the following CGS update equation:

$$\mathbb{P}(z_{m,n,r}=k\mid z_{-(m,n,r)},w,\alpha,\beta)\propto(\boldsymbol{C}_{k,m,*,*}^{-(m,n,r)}+\alpha_k)\times\frac{(\boldsymbol{C}_{k,*,n,v}^{-(m,n,r)}+\beta_{n,v})}{(\boldsymbol{C}_{k,*,n,*}^{-(m,n,r)}+\beta_{n,*})}\qquad(\text{here, }w_{m,n,r}=v) \tag{5}$$

Once the strain identifier $z_{m,n,r}$ is sampled using Equation 5 above, its sampled value is used to update the counts $\boldsymbol{C}_{k,m,*,*}$ and $\boldsymbol{C}_{k,*,n,v}$ (recall $w_{m,n,r}=v$).

The unnormalized $\boldsymbol{\theta}$ and $\boldsymbol{\phi}$ can then be determined from the above updated counts as:

$$\boldsymbol{\theta}_{m,k}=\boldsymbol{C}_{k,m,*,*}$$
$$\boldsymbol{\phi}_{k,n,q}=\boldsymbol{C}_{k,*,n,q}$$

When calculating KL divergence measures that involve inferred $\boldsymbol{\phi}$ distributions, we add a pseudocount of $\beta_{n,q}$ to the estimate of $\boldsymbol{\phi}_{k,n,q}$ above to avoid infinite KL divergence values.

#### 1.2. From a reference database to global/sample-specific dictionaries

The reference database of Demixer is constructed using the 7684 barcoding SNP-alleles identified by Napier et al. [8] to distinguish 91 TB lineages and sublineages (all of which are referred to simply as strains for convenience; the SNP-alleles in the reference database are also called as reference SNP-alleles). As mentioned in the main text, we have access to a lineage tree that represents the hierarchical (ancestor-descendant) relationship between all the strains (lineages/sublineages) in the reference database. We now derive two types of databases, RDB1 and RDB2.

RDB1 includes only the unique SNP-alleles of each strain - a SNP-allele is called unique for a strain $j$ if it is present in that strain and additionally absent in all other non-descendant strains of strain $j$ in the reference database. Note that these unique SNP-alleles exclusively identify a strain (excepting all its descendants). RDB1 can be viewed as a binary matrix of dimension $7684\times L$, where L is the number of strains. Note that, more the number of unique SNP-alleles for a specific strain in RDB1, the more information we've on this strain in the reference database, making it less likely for Demixer to miss the identification of that strain.

RDB2 consists of all SNP-alleles recorded in the reference database (including both unique and non-unique SNP-alleles of strains). It is also represented as a matrix and its entry RDB2$[(n,v),j]=1$ if SNP-allele $(n,v)$ is present in strain j or any ancestor of strain $j$. In other words, the SNP-alleles of each strain is propagated down to all its descendant strains; with the assumption being that a SNP-allele in a strain is also present in all its children and descendant strains (for example, if A100T belongs to strain 1.2.1, then A100T will also belong to all its descendants 1.2.1.x). Using these RDB* databases, we derive the dictionaries GD and SD$_m$; please refer Algorithm 1 for the usage of these dictionaries in the parallelized CGS algorithm.

To construct the global dictionary (GD), we start with the pruning of the lineage tree. The rows corresponding to SNP-alleles that are absent in all input samples are removed from RDB1, followed by the removal of columns/strains with less than 5 non-zero entries - the SNP-alleles and strains removed during this process are also removed from RDB2, and the resulting RDB2' is used to create the global dictionary (GD). In this dictionary, each key corresponds to the SNP-allele $(n,v)$, and the related value is the list of all strains $j$ such that RDB2'$[(n,v),j]$=1.

To construct the sample-specific dictionary (SD$_m$), we follow the same procedure as in the above paragraph, but we start with the removal of SNP-alleles that are absent in sample $m$ alone (instead of those absent in all input samples); and also remove only the columns/strains with 0 non-zero entries (instead of less than 5 non-zero entries as before).

#### 1.3. Postprocessing/interpretation steps

The different steps involved in postprocessing the strains inferred by Demixer are described in this section.

**Mapping inferred to reference strains:** Given Demixer's output, we would like to know if any of the inferred strains output by Demixer can be mapped to a known reference strain (whose unique mutations/SNP-alleles are in a database called RDB1; see Supplementary Section 1.2). For each inferred strain $q$ ($1 \leq q \leq K$), we check if it is the same as a particular reference strain $p$ by first computing a dissimilarity measure (Kullback–Leibler or KL divergence $D_{KL}$ [3] of the inferred distribution $\boldsymbol{\phi}_q$ from the expected reference strain distribution $\boldsymbol{\phi}_p^{\text{ref}}$) as given in equation 6 below. Let $N_p$ be the set of SNPs such that one of its alleles uniquely identifies the reference strain $p$, i.e., $N_p = \{n : \left(\sum_{v \in \{A,C,G,T\}} \text{RDB1}[(n,v),p]\right) = 1\}$. Then,

$$D_{(KL)}(p \mid\mid q) := \frac{1}{|N_p|} \sum_{n \in N_p} D_{KL}(\boldsymbol{\phi}_{p,n}^{\text{ref}} \mid\mid \boldsymbol{\phi}_{q,n}) \tag{6}$$

where the Strain-SNP-allele distributions $\boldsymbol{\phi}_{p,n}^{\text{ref}}$ is given by the four-element vector $\text{RDB1}[(n,v),p]_{v \in \{A,C,G,T\}}$, and $\boldsymbol{\phi}_{q,n}$ is as estimated by Demixer.

The inferred strain $q$ is assigned to the reference strain for which the KL divergence value is minimum and at most 1.5. If there are multiple such strains, we map $q$ to the reference strain that is deepest in the hierarchical tree relating these strains (and finally if ties still remain, which may happen rarely, we break the ties arbitrarily). If the KL divergence is more than 1.5 for all reference strains, then strain $q$ is called as a *de novo* or unmapped strain. Note that any inferred strain whose proportion is 0 in all the samples is excluded from all post-processing analyses.

Note that the above procedure maps each inferred strain to zero or one reference strain, but a reference strain could be mapped to 0, 1, or more inferred strains. The proportion of a reference strain mapping to more than one inferred strains is simply the sum of the estimated Sample-Strain proportions of the corresponding inferred strains.

**Fine-tuning the mapped strains using the lineage tree:** Complex datasets (like real-world datasets or simulated datasets mimicking real-world scenarios) may cause our Demixer to call a sample that contains only a single lineage (pure) as a mixture of different strains/sub-lineages within the single lineage (mixed infection). This may happen if additional mutations of the lineage missing from the reference database are present in the sample. To mitigate this issue, we perform sample-level post-processing for a dataset of interest using the lineage tree, which is a hierarchical tree relating all lineages/sub-lineages in the reference database (derived using naming conventions; e.g., 1.2.1 is a sub-lineage of 1.2). For each sample, we consider the set of reference strains mapped to at least one inferred strain in the sample, and process these reference strains one pair at a time in a certain fixed ordering. If a reference strain pair has an ancestor-descendant relation in the lineage tree, then we will retain only the tree node with the larger estimated proportion and add to it the smaller proportion of the other node. For instance, if the inferred strains of sample $m$ gets mapped to reference lineages 1.2 ($p$) and 1.2.1 ($p'$), then we retain only lineage 1.2.1 if $\boldsymbol{\theta}_{m,p'} > \boldsymbol{\theta}_{m,p}$ and only 1.2 otherwise. After processing all such reference strain pairs, the set of retained strains along with the unmapped (*de novo*) inferred strains if any constitute the final strains called for the sample. A sample with only one final strain is reported as non-mixed/pure by Demixer (see Algorithm 2 in Supplementary Information for detailed steps of the post-processing heuristic). Please note that in the application of Demixer to all samples analyzed in this work, we detected either 0 or 1 *de novo* strains per sample; with the exception of a single sample (in the CRyPTIC dataset) where two *de novo* strains were detected – for simplicity, we merged both these strains into a single *de novo* strain by adding up their proportions as above.

**Quality checks using SNP plots/modes and *de novo* filtering:** Complex/noisy datasets also calls for the use of additional SNP-based quality checks and filters during post-processing. For each sample $m$ in a dataset of interest, we plot the histogram of minor allele proportions of heterozygous SNPs. The x-axis of this histogram plot (also called the SNP plot) corresponds to the minor allele proportions of the heterozygous SNPs and the y-axis corresponds to the frequency (count) of such SNPs. Note that a SNP $n$ is heterozygous in sample $m$ if it has reads supporting at least two alleles, i.e., if $\sum_v \mathbb{1}_{\{\boldsymbol{S}_{m,n,v} > 0\}} \geq 2$. Also, if a SNP has more than one minor allele, we sum up all these alleles' proportions to get the minor allele proportion for this plot. From the SNP or histogram plot of sample $m$, we can inspect the highest frequency histogram bin to estimate the mode of the minor allele proportions of heterozygous SNPs (denoted $o_m$; see Algorithm 2 for details) as well as its corresponding frequency (the number of SNPs, denoted $f_m$).

Given a dataset, we inspect the distribution of $f_m$ values across all samples in the dataset, and use it to assign a confidence measure to each specific sample identified as mixed by Demixer. Let cutoff1 be the 25th and cutoff2 the 75th percentile of $\{f_m\}_{m=1}^M$; note that these cutoffs are expressed in terms of the number of SNPs. Then for a specific sample $m$, we apply the thresholds $f_m <$ cutoff1, cutoff1 $\leq f_m \leq$ cutoff2, and $f_m >$ cutoff2 to categorize Demixer's prediction for the sample as low, medium, and high confidence call respectively. Additionally, if Demixer's prediction for the sample (after the post-processing steps above) results in more than one inferred strains, with one mapped to a reference strain and the rest being *de novo* strain(s), we report the sample as mixed only if it's a high confidence call; otherwise, we remove (filter out) the *de novo* strain(s) and thereby report the sample as non-mixed with only the single reference strain.

These confidence cutoffs and *de novo* filtering will influence different downstream analyses. For $F1$ score calculation, the *de novo* filtering step will have an effect, especially for low-coverage benchmark datasets. For the Malawi dataset, we use these cutoffs to compare the performance of Demixer with an existing work MixInfect. For the CRyPTIC dataset, we apply similar thresholds ($f_m >$ cutoff2 and $o_m > 0.05$) to call a mixed sample detected by Demixer as high confidence, and thereafter test for association between drug resistance and mixed infection. Note that the $f_m$ and $o_m$ thresholds respectively ensure a sufficient number of SNPs and reads support the minority strain.

**Hierarchical clustering of the inferred strains, and *in vitro* dataset processing:** We compute the KL divergence measure between the inferred Strain-SNP-allele distribution of each strain $q'$ from every other strain $q > q'$ using the equation below, resulting in a lower-triangular KL matrix of size $K \times K$. This matrix is given as input to the hierarchical clustering algorithm (implemented using hclust function of R [11] stats package) to obtain the tree structure of the inferred strains. The nodes in the tree captures the lineages/sub-lineages of the inferred strains.

$$D_{(KL)}(q \mid\mid q') := \frac{1}{N} \sum_{n=1}^{N} D_{KL}(\phi_{q,n} \mid\mid \phi_{q',n})$$

For the *in vitro* dataset, since the ground-truth is given in terms of majority and minority proportions, we aggregated the inferred strains in each sample into at most two strains by clustering the closer strains. Specifically, for each sample we clustered the strains using the above tree structure at different heights, starting from the leaves and until two clusters remain (using the cutree function of R). This procedure results in one clustered strain or two clustered strains called for each sample. The sum of the proportions of strains in each cluster is then taken as the majority/minority proportions present in the sample.

We perform two types of evaluation using these calls on the *in vitro* dataset. In proportion-based evaluation, the true and estimated proportions are plotted and compared with each other, regardless of the clustered strain identities (in order to facilitate comparison of Demixer with methods which doesn't infer strain identities like SplitStrains). In strain-identity-based evaluation, we inspect if the lineage of the clustered strains (i.e., the consensus lineage of all the inferred strains within the cluster) is the same as the ground-truth lineage. The lineage of an inferred strain is obtained as before (using the KL divergence based mapping procedure above, refined using the above clustering procedure instead of the fine-tuning procedure using the lineage tree).

#### 1.4. Datasets

This section briefly describes the different synthetic and real-world datasets that we have employed in our study to understand and validate our Demixer method's performance.

**LDAmix1 and LDAmix2 datasets:** These are synthetic datasets created based on the modified LDA's generative process used in the Demixer method. Dataset LDAmix1 consists of 100 samples, and each sample is composed of 3 mixed strains. Each strain consists of 50 distinct SNPs. The proportion of the majority strain (strain 1) is drawn from the Gaussian distribution $\mathrm{N}(0.6, 0.05)$ with a mean 0.6 and standard deviation 0.05, that of the minority strain (strain 2) from another Gaussian $\mathrm{N}(0.2, 1)$ with a mean 0.2 and standard deviation 1, and the remaining proportion pertains to strain 3. LDAmix2 consists of 100 samples with 4 mixed strains, each with 50 unique SNPs. The minority strain (strain 4) is present only in 10% of the samples in order to determine the ability of Demixer to detect the strains present in a deficient proportion in fewer samples. The majority strain (strain 1) is drawn from $\mathrm{N}(0.6, 0.05)$, strain 2 from $\mathrm{N}(0.2, 1)$, strain 4 from $\mathrm{N}(0.1, 0.05)$, and strain 3 constitutes the remaining proportion. Each sample in both of these datasets consists of "n (no.of strains) $\times$ 50" SNPs with 100 reads for each SNP. Please note that we could also have used Dirichlet instead of Gaussian distributions for sampling the strain proportions above.

**ART-TBmix1 and ART-TBmix2 datasets:** Each ART-TBmix* dataset consists of 800 samples at four different coverages: 10x, 20x, 90x-10x, and 70x-30x. Each sample is combined with 4 strains in 10x and 20x subsets while in 90x-10x, and 70x-30x subsets, it is mixed with 2 strains. The strains for the datasets ART-TBmix1 and ART-TBmix2 are selected from QuantTB's [1] database consisting of 2165 strains and Napier's [8] work consisting of 89 strains, respectively. The representative strains in both these datasets are generated by replacing the SNPs related to that strain in the reference H37Rv genome. The WGS reads for each sample are generated using the ART simulator (Version 2.5.8) with the default settings for the Illumina HiSeq 2500 platform, at a read length of 101 bp, per base sequence quality scores 20 to 30 and the quality scores are shifted by 9 to introduce sequencing errors as described in [1] and [6]. The coverage option is set accordingly for generating strains at different coverages 10x, 20, 70x, 30x, and 90x respectively for the different subsets. The four-strain sample is generated by selecting four strains at random from the respective databases and paired-end WGS reads for the individual strains are simulated at the specific coverage level. The reads of all four strains are then combined to generate the mixed sample. In the same manner, two strains are generated at two different coverages (for instance, 70x and 30x), to synthesize a sample mixed with two strains at 70x-30x coverage.

**ARTmix dataset:** ARTmix has been generated to determine whether Demixer could delineate the strains accurately under different scenarios such as when all the references are known, and when a few references are known. ARTmix dataset consists of 50 samples to emulate the real-world conditions where various samples exhibit diverse combinations of mixed strains. Each sample is either a single strain or a mixed strain composed of 2 or 3 strains. Each strain in the sample is selected at random from a reference set of 7 strains. The reference strains are obtained by replacing nearly 100 SNPs in the reference H37Rv genome and the WGS data is generated as per the settings described in SplitStrains [6] work using the ART simulator. The process of determining whether a sample contains one, two, or three strains is based on a random selection from a $\mathrm{Uniform}(1, 4)$ distribution. In the case of two strain mixed samples, the majority strain (strain 1) is drawn from $\mathrm{N}(a, 1)$, such that $40 \leq a \leq 70$ is a random integer and strain 2 constitutes the remaining proportion. Similarly, for the three strain mixed samples, majority strain (strain 1) is drawn from $\mathrm{N}(a, 1)$, such that $60 \leq a \leq 70$ is a random integer, strain 2 is drawn from $\mathrm{N}(b, 1)$, such that $0 \leq b \leq 20$ is a random integer and strain 3 constitutes the remaining proportion.

**Malawi and CRyPTIC datasets:** We ran Demixer on two real-world TB Datasets namely Malawi and CRyPTIC to identify mixed infection. The Malawi TB isolates are collected from patients in Malawi, a Southeastern African region having higher rates of TB and HIV co-infection [7]. Samples were collected from TB patients at regular intervals during the period 1996-2010 to understand whether the prevalence of HIV has any role in reinfection/relapse of TB. We have also analyzed 1963 isolates (downloaded from the publicly available datasets with accession numbers PRJEB2794 and PRJEB2358) as in [12]. CRyPTIC is a collaborative effort that is focused on the better identification of drug-resistant TB by analyzing the WGS data. They work with TB research institutions from 27 countries worldwide and have collected WGS data of nearly 12289 isolates along with their responses to 13 tuberculosis drugs [4][5]. We used 12287 isolates that we were able to download in a proper vcf format in our analysis.

#### 1.5. Model selection and hyperparameter tuning

Symmetric or asymmetric priors can be chosen for the hyperparameters $\alpha$ and $\beta$ of the LDA model. In symmetric prior, the assumption is that each strain is equally likely in a sample and each SNP-allele is equally likely in a strain, whereas asymmetric prior allows certain strain/SNP-alleles to occur more often than others in a set of samples. Though symmetric priors are commonly chosen, Wallach et al. [14] have shown the advantages of choosing asymmetric Dirichlet prior over the document-topic distribution, whereas there is limited improvement in choosing asymmetric prior over word-topic distribution. A similar hypothesis has been put forward in [13] by testing different combinations of symmetric and asymmetric priors over $\boldsymbol{\phi}$ and $\boldsymbol{\theta}$ distributions using coherence score and interpretability measures. In [9], topic-dependent smoothing coefficients for words were employed to facilitate the identification of topics with incoherent words, rather than relying on an asymmetric prior for topic-word distribution. We follow a similar method here.

Specifically, we employed 5 different hyperparameter combinations $(1, 0.01)$, $(0.01, A)$, $(0.01, B)$, $(A, 0.01)$ and $(B, 0.01)$ (including asymmetric prior for $\alpha$ parameter and topic smoothing coefficients for $\beta$ parameter) to identify the default hyperparameters for the model. The first element in each set corresponds to the $\alpha$ hyperparameter and the second element to the $\beta$ hyperparameter. $(1, 0.01)$ is the simple prior chosen assuming that a sample contains only a few minority strains and either the reference or alternate allele occurs with higher probability at a SNP position. For the asymmetric combination, if the number of strains is assumed to be 3, A will take the values [0.01, 1, 1] and B [0.01, 10, 1]. Here, 0.01 indicates the reference strain for which the SNP-alleles are known, and the remaining values correspond to the *de novo* strains. A is assigned values assuming that the minority strains are equally likely to occur, whereas, in B, the assumption is that among the minority strains, one would occur in the majority than the other. Similarly, for 4 strain cases, A and B would correspond to [0.01, 1, 1, 1] and [0.01, 10, 1, 1] respectively.

#### 1.6. Comparison of Demixer and traditional LDA

One of the key differences between our proposed Demixer and traditional text LDA is that instead of having a single $\boldsymbol{\phi}$ matrix for all the SNP-allele combinations, we have a $\boldsymbol{\phi}$ matrix for each SNP. To empirically understand this difference, we applied traditional LDA and Demixer in *de novo* mode (no reference database given) on a special dataset where samples have strains mixed in roughly equal proportions. Note that the inference of mixed strains' identities/proportions in this dataset is difficult due to the equal proportions of strains. This dataset specifically comprised 53 samples (2 pure and 51 mixed with 2 synthetic strains) such that the proportion of the majority strain ranges from 49 to 100% in 1% increment in the mixed samples. Each synthetic strain is obtained by replacing 100 random SNPs in the reference H37Rv genome. We used Python Gensim library for the traditional LDA modeling [10]. In this implementation, each read that supports a SNP-allele combination is represented as a word in a document, and then fed to the LDA model. The traditional LDA neither directly captures the sequencing depth (i.e., the number of reads mapping to a SNP) nor the distribution of alleles within each SNP, whereas this aspect is inherently built within the framework of our SNP-LDA model. The proportion of one of the strains determined by traditional text LDA and Demixer on the samples in this dataset was compared with its actual proportion. Traditional LDA was unable to determine the proportions accurately for the samples with equal proportions of strains. The ratios determined by Demixer were very close to the actual ratios, as illustrated in Figure S2, indicating that our proposed modeling approach determines the proportions with high accuracy.

#### 1.7. Implementation details

All the experiments are conducted on an Intel(R) Xeon(R) Platinum 8180 CPU @ 2.50GHz processor running CentOS with 112 cores and 1 TB RAM. The code for preprocessing the input vcf file is implemented in Python. The tools from Python *scikit-allel* framework are used for reading and processing the vcf files. The parallelization of the CGS algorithm is implemented using the OpenMP library. The gsl_ran_multinomial function from GNU Scientific Library (GSL) is utilized to generate random samples from a multinomial distribution in the CGS algorithm. The postprocessing steps of Demixer are implemented again in Python using the estimated model parameters of the SNP-LDA model. For simplicity, we have reduced the 3D matrices $\boldsymbol{S}$ and $\boldsymbol{\phi}$ to 2D during the code development of Demixer. We used the programming language and statistical software environment R to perform hierarchical clustering of the strains detected in the *in vitro* dataset samples.

#### 1.8. Using Demixer for analysing new sample(s)

Instead of learning Demixer separately on each dataset, we can also run Demixer in a training+testing mode, wherein the Demixer model learnt from a large diverse training dataset (CRyPTIC in our case) can be used to inspect the strains in a small test dataset containing one or more new samples. To facilitate this mode, Demixer is applied on the training dataset in an unsupervised (without class labels) fashion to learn the model parameters $\boldsymbol{\theta}^{\text{train}}$ and $\boldsymbol{\phi}^{\text{train}}$; of these, we utilize only the $\boldsymbol{\phi}^{\text{train}}$ parameter to learn the test samples' strain proportions $\boldsymbol{\theta}^{\text{test}}$. The test dataset goes through the same three steps of Demixer, preprocessing, estimating $\boldsymbol{\theta}^{\text{test}}$, and post-processing, but with a few key changes. In preprocessing, we ensure the same set of SNPs are analyzed in both training and test datasets (by using the force call option in FreeBayes tool with the merged multisample .vcf file of the training data as its input). When estimating $\boldsymbol{\theta}^{\text{test}}$, we initialize the strains of reads using the critical information in $\boldsymbol{\phi}^{\text{train}}$ parameter as shown in equation 7 below, and run the rest of the hybrid SNP-LDA parallelized CGS algorithm as is for a fixed number of iterations (100 for single sample and 500 for a batch of samples to update $\boldsymbol{\theta}^{\text{test}}$ and $\boldsymbol{\phi}^{\text{test}}$, and eventually report the former and ignore the latter). Note that the CGS update formula (equation 5) is used as is for the CGS iterations, but its revised form (equation 7 below that uses $\boldsymbol{\phi}^{\text{train}}$ and assumes $\boldsymbol{\theta}$ is based only on its prior) is used for initialization alone.

$$\begin{aligned}\mathbb{P}(z_{m,n,r} = k \mid w_{m,n,r} = v, \boldsymbol{\phi} = \boldsymbol{\phi}^{\text{train}}, \boldsymbol{\theta} = \{\alpha_k\}) \\ \propto \alpha_k \cdot \frac{(\boldsymbol{wt}_{n,v} \boldsymbol{\phi}^{\text{train}}_{k,n,v} + \beta_{n,v})}{(\sum_{v'} \boldsymbol{wt}_{n,v'} \boldsymbol{\phi}^{\text{train}}_{k,n,v'} + \beta_{n,v'})}\end{aligned} \tag{7}$$

For this training+testing mode to work, the training dataset should be reasonably large and diverse. Specifically, a diverse set of strains (established reference strains as well as newly discovered strains) should be represented in a sufficiently large number of samples in the training dataset. Large training datasets with sufficient sequencing depth can also allow us to mitigate errors if any in the mutations recorded in the reference database. Based on these factors and our analysis of applying Demixer on different datasets of sizes ranging from 48 to 12,287, we propose a heuristic that the training dataset must have at least 100 samples when learning a Demixer model in the training+testing mode.

#### 2. Supplementary Algorithms

**Algorithm 1 Parallel Collapsed Gibbs Sampling:** This pseudocode pertains to Demixer's default mode known as hybrid SNP-LDA ( to obtain non-hybrid SNP-LDA variants, the dictionaries $\mathrm{SD}_{m=1}^{M}$ or GD (see Supplementary Section 1.2 for details of SD and GD) that map known SNP-alleles to the strains they are present should be emptied out before being provided as input). The algorithm begins by assigning strains to each read of a sample either at random or using reference SNP-alleles. During each CGS iteration, strains to each read of a sample are reassigned based on $\boldsymbol{\theta}$ and $\boldsymbol{\phi}$, followed by the updation of these matrices.

**Input:** Sample-SNP-allele matrix $\boldsymbol{S}_{M\times N\times 4}$, SNP weights $\boldsymbol{wt}_{N\times 4}$, the sample-specific dictionaries $\mathrm{SD}_{m=1}^{M}$, and global dictionary GD (with $SD_m[j]$ or $GD[j]$ mapping SNP-allele $j$ to the set of strains present in the $m$-th sample or across all samples respectively).
**Output:** Inferred model parameters – Sample-Strain matrix $\boldsymbol{\theta}_{M\times K}$, Strain-SNP-allele matrix $\boldsymbol{\phi}_{K\times N\times 4}$.

$K \leftarrow K' + 2$ (determined as per the heuristic for choosing $K$)
Initialize $\boldsymbol{\theta}$, $\boldsymbol{\phi}$, $\boldsymbol{C}_{K\times M\times N\times 4}$ to zeros
\# pragma omp parallel for
**for** $i = 1, 2, \ldots, M$ **do**
  **for** $j = 1, 2, \ldots, N$ **do**
    **for** $v = 1, \ldots, 4$ **do**
      **if** $(j, v)$ in $\mathrm{SD}_i$ **then**
        u $\leftarrow$ prob($\mathrm{SD}_i[(j, v)]$)
      **else**
        u $\leftarrow [\frac{1}{K}, \frac{1}{K}, \ldots, \frac{1}{K}]$ ($K$-length vector)
      **end if**
      $\boldsymbol{C}_{1:K,i,j,v} \leftarrow$ Multinomial($\boldsymbol{S}_{ijv}$, u)
      Update $\boldsymbol{\theta}_{i,1\ldots K}$ using $\boldsymbol{C}_{1:K,i,*,*}$
    **end for**
  **end for**
**end for**
Update $\boldsymbol{\phi}$ using $C$
**for** $iter = 1, 2, \ldots,$ max_iterations **do**
  \#pragma omp parallel for
  **for** $i = 1, 2, \ldots, M$ **do**
    **for** $j = 1, 2, \ldots, N$ **do**
      $\mathrm{a}_{i,1:K} \leftarrow \boldsymbol{\theta}_{i,1:K}$
      $\mathrm{b}_{1:K,j,1:4} \leftarrow \boldsymbol{wt}_{j,1:4} * \boldsymbol{\phi}_{1:K,j,1:4}$
      $\boldsymbol{\theta}_{i,1:K} \leftarrow \boldsymbol{\theta}_{i,1:K} - (\boldsymbol{wt}_{j,1:4} * \boldsymbol{C}_{1:K,i,j,1:4})$
      **for** $v = 1, \ldots, 4$ **do**
        **if** $(j, v)$ in GD **then**
          strains $\leftarrow$ GD[$(j, v)$]
          u $\leftarrow$ a[strains] .* b[strains]
        **else**
          u $\leftarrow \mathrm{a}_{1:K}$ .* $\mathrm{b}_{1:K}$
        **end if**
        $\boldsymbol{C}_{1:K,i,j,v} \leftarrow$ Multinomial($\boldsymbol{S}_{ijv}$,u)
        Update $\boldsymbol{\theta}_{i,1:K}$ using $\boldsymbol{C}_{1:K,i,*,*}$
      **end for**
    **end for**
  **end for**
  Update $\boldsymbol{\phi}$ using $C$
**end for**
**for** $m = 1, 2, \ldots, M$ **do**
  **for** $k = 1, 2, \ldots, K$ **do**
    $\boldsymbol{\theta}_{m,k} = \frac{\boldsymbol{C}_{k,m,*,*}}{\boldsymbol{C}_{*,m,*,*}}$
  **end for**
**end for**
**for** $k = 1, 2, \ldots, K$ **do**
  **for** $n = 1, 2, \ldots, N$ **do**
    **for** $v = 1, \ldots, 4$ **do**
      $\boldsymbol{\phi}_{k,n,v} = \frac{\boldsymbol{C}_{k,*,n,v}}{\boldsymbol{C}_{k,*,n,*}}$
    **end for**
  **end for**
**end for**

- prob($\mathrm{SD}_i[(j, v)]$): If $\mathrm{SD}_i[(j, v)]$ returns more than 1 strain, then the strain at a deeper level in the hierarchy will be assigned a higher probability (0.95) than the other strains. In case of ties, the strain that is seen first during processing will be assigned higher probability.
- .* indicates element-wise multiplication.

**Algorithm 2 Post-processing steps of Demixer:** A brief overview of the postprocessing steps and the order in which the different postprocessing steps are done by Demixer is shown here. Please refer Supplementary Section 1.3 for a detailed description of each postprocessing step.

**Input:** Inferred model parameters, $\boldsymbol{\theta}$, $\boldsymbol{\phi}$, output by Algorithm 1
**Output:** Phylogeny to visualize the inferred strains; Final Proportions $\boldsymbol{FP}_{M\times L}$, Final Strains $\boldsymbol{FS}_{M\times L}$, and (Mixed/Non-mixed) Assignments $\boldsymbol{AS}_M$
**Note:** $L$ indicates the maximum number of strains that could be present in a sample after postprocessing/merging. Also, $\boldsymbol{FP}_{M\times L}$ stores the final refined proportions, $\boldsymbol{FS}_{M\times L}$ the final strains, and $\boldsymbol{AS}_M$ the mixed vs. non-mixed assignment for each of the $M$ samples (after postprocessing/merging).

/* **Hierarchical clustering of the inferred strains:** */
Compute and output a phylogeny relating all the inferred strains, excepting strains whose estimated proportion is zero across all samples (R hclust function is used as described in Supplementary Section 1.3).

/* **Mapping inferred to reference strains:** */
Setup mapped_reference_strainid$_K$ array as follows. Map each inferred strain $k$ to the closest reference strain $p$ with KL-divergence at most 1.5 (as described in Supplementary Section 1.3), and store this information as "mapped_reference_strainid[$k$] = $p$". If no such reference strain exists, then store "mapped_reference_strainid[$k$] = concat('*de novo* strain',$k$) ".

**for** $i = 1, 2, \dots, M$ **do**
    prop_list ← non-zero-values($\boldsymbol{\theta}_m$)    (get the non-zero strain proportions)
    id ← mapped_reference_strainid[non-zero-indices($\boldsymbol{\theta}_m$)] (get the id's of mapped reference strains with non-zero proportion)
    id ← sort(id); prop_list also sorted using same ordering; (sorting required for prefix check below)
    **if** len(id) = 1 **then**
        $\boldsymbol{AS}_m$ ← "Non-mix"; $\boldsymbol{FS}_m$ ← id; $\boldsymbol{FP}_m$ ← prop_list;
        **continue** to next sample
    **end if**

    /* **Fine-tuning (aka merging) the mapped strains using the lineage tree:** */
    **for** $j = 1, 2, \dots, len(\text{id}) - 1$ **do**
        /* Prefix condition checks if strain j (e.g., lineage 4.2) is ancestor of strain j+1 (e.g., lineage 4.2.1) in the lineage tree */
        **if** id$_j$ = prefix(id$_{j+1}$) and ids $j, j+1$ are not *de novo* strains **then**
            **if** prop_list$_j$ > prop_list$_{j+1}$ **then**
                id$_{j+1}$ ← id$_j$    (Assign the id of the ancestor to the merged strain)
            **end if**
            prop_list$_{j+1}$ ← prop_list$_j$ + prop_list$_{j+1}$    (Merge)
            Flag strain $j$ as "removed".
        **else**
            Do not merge
        **end if**
    **end for**
    $\boldsymbol{FS}_m$ ← ids of all strains NOT flagged as "removed" above; $\boldsymbol{FP}_m$ ← corresponding entries of prop_list;

    /* Special case handling: **Quality checks using SNP plots/modes and *de novo* filtering:** */
    prop$_m$ ← $[ma_1, ma_2, \dots, ma_H]$    ($ma_h$ is the minor allele proportion at $h$-th heterozygous SNP out of all $H$ heterozygous SNPs in the sample)
    $f_m$ ← frequency of the most-frequent bin in histogram(prop$_m$)
    $o_m$ ← mode of prop$_m$ (estimated using the mode of $ma_h$ values falling within the most-frequent histogram bin)
    /* Note: "$f_m >$ cutoff2" is a high confidence call; see Supplementary Section 1.3 for details */
    **if** $\boldsymbol{FS}_m$ contains one reference strain and at least one *de novo* strain AND $f_m \leq$ cutoff2 **then**
        Remove (filter out) all *de novo* strain(s) from $\boldsymbol{FS}_m$, and update $\boldsymbol{FP}_m$ accordingly
    **end if**

    **if** len($\boldsymbol{FS}_m$) = 1 **then**
        $\boldsymbol{AS}_m$ ← "Non-mix"
    **else**
        $\boldsymbol{AS}_m$ ← "Mix"
    **end if**
**end for**

#### 3. Supplementary Tables

**Table S1. Summary of the datasets used in the study**

| Dataset | Number of Strains | Synthetic/ Real World | Number of Samples |
|---|---|---|---|
| LDAmix1 | 3 | Synthetic - Demixer's Generative Process | 100 (all mixed) |
| LDAmix2 | 4 | Synthetic - Demixer's Generative Process | 100 (all mixed) |
| ART-TBmix1 | 2,4 | Synthetic - Art Simulator | 800 (all mixed) |
| ART-TBmix2 | 2,4 | Synthetic - Art Simulator | 800 (all mixed) |
| ARTmix | 1,2,3 | Synthetic - Art Simulator | 50 (18 pure & 32 mixed) |
| Malawi | unknown | Real-world Dataset | 1963 |
| CRyPTIC | unknown | Real-world Dataset | 12287 |

**Table S2. Empirical estimation of the constant $D$ in Gibbs sampler's weighted update equation:** Demixer is run on the *in vitro* dataset with different values for $D$ ranging from 1 to 10 (100 runs for each value of $D$) to determine its optimal value in the update equation 3 (and the related $\boldsymbol{wt}_{n,v}$ formula) in the main text. The reported values are the average relative errors and standard deviations computed across 100 runs for each value of D.

| $D$ | 0 (no weight) | 1 | 2 | 3 | 4 | 5 | 6 | 7 | 8 | 9 | 10 |
|---|---|---|---|---|---|---|---|---|---|---|---|
| **Weight ($\boldsymbol{wt}_{n,v}$)** | 1 | 2 | 2 | 3 | 4 | 4 | 5 | 6 | 7 | 7 | 8 |
| **Average Relative Error** | 0.5432 | 0.5166 | 0.5589 | **0.2920** | 0.3413 | 0.3407 | 0.3866 | 0.4114 | 0.4282 | 0.4169 | 0.3897 |
| **Standard deviation** | 0.2157 | 0.2092 | 0.1950 | **0.0052** | 0.0216 | 0.0220 | 0.0449 | 0.0918 | 0.1369 | 0.1445 | 0.1860 |

**Table S3. Weights estimated for different datasets:** The weight assigned to the reference alleles in each of the synthetic and real-world datasets determined as per the $\boldsymbol{wt}_{n,v}$ formula described in the main text is shown. As mentioned in the main text, $N'$, $A$, $K$, and $D$ represent the number of SNPs, the average number of reference SNP-alleles per sample, the number of strains, and the constant respectively.

| Dataset | Type | $N'$ | $A$ | $K$ | $D$ | $\boldsymbol{wt}_{n,v}$ | adjusted $\boldsymbol{wt}_{n,v}$ |
|---|---|---|---|---|---|---|---|
| **LDAmix1 (synthetic)** | **non-hybrid** | 150 | 0 | 3 | 3 | 0 | 1 |
| | **hybrid** | 150 | 50 | 3 | 3 | 2 | 2 |
| **LDAmix2 (synthetic)** | **non-hybrid** | 200 | 0 | 4 | 3 | 0 | 1 |
| | **hybrid** | 200 | 50 | 4 | 3 | 3 | 3 |
| **ART-TBmix1 (synthetic)** | **10x** | 359372 | 2203 | 786 | 3 | 1 | 2 |
| | **20x** | 95319 | 2321 | 672 | 3 | 1 | 2 |
| | **70x-30x** | 60034 | 1434 | 365 | 3 | 1 | 2 |
| | **90x-10x** | 59081 | 1396 | 365 | 3 | 1 | 2 |
| **ART-TBmix1 (synthetic)** | **10x** | 281228 | 439 | 83 | 3 | 24 | 24 |
| | **20x** | 6676 | 447 | 82 | 3 | 1 | 2 |
| | **70x-30x** | 5543 | 233 | 82 | 3 | 1 | 2 |
| | **90x-10x** | 5539 | 230 | 82 | 3 | 1 | 2 |
| **ARTmix (synthetic)** | **all references known** | 729 | 164 | 7 | 3 | 1 | 2 |
| | ***de novo* mode** | 729 | 111 | 7 | 3 | 3 | 3 |
| ***in vitro*** | **benchmark** | 4818 | 313 | 19 | 3 | 3 | 3 |
| **Malawi** | **real-world** | 15625 | 246 | 43 | 3 | 5 | 5 |
| **CRyPTIC** | **real-world** | 74495 | 131 | 71 | 3 | 24 | 24 |

**Table S4. Time and space usage of Demixer for the experimental/real-world datasets**: The time and space used by Demixer for preprocessing and running parallel CGS algorithm for 1000 iterations is shown. The memory usage reported here is the maximum virtual memory utilized as determined by the Unix top command.

| **Dataset** | **Run-time** | | **Memory** | |
|---|---|---|---|---|
| | **Pre-processing** | **Parallel CGS** | **Pre-processing** | **Parallel CGS** |
| *in vitro* | 11.4 seconds | 54 seconds | 0.16 GB | 4.067 GB |
| Malawi | 3.57 minutes | 20.41 minutes | 1.61 GB | 14.3 GB |
| CRyPTIC | 1.62 hours | 17.68 hours | 46.51 GB | 368.1 GB |

**Table S5. Precision, Recall and $F1$ scores of Demixer and QuantTB on ART-TBmix1 dataset**:

| **Coverage** | **Demixer** | | | **QuantTB** | | |
|---|---|---|---|---|---|---|
| | **Precision** | **Recall** | **$F1$ score** | **Precision** | **Recall** | **$F1$ score** |
| **10x** | 0.91 | 0.91 | 0.91 | 0.98 | 1.00 | 0.99 |
| **20x** | 1.00 | 0.99 | 0.99 | 0.98 | 1.00 | 0.99 |
| **90x-10x** | 0.99 | 0.99 | 0.99 | 0.99 | 1.00 | 0.99 |
| **70x-30x** | 1.00 | 1.00 | 1.00 | 0.99 | 1.00 | 1.00 |

**Table S6. Precision, Recall and $F1$ scores of Demixer and QuantTB on ART-TBmix2 dataset**:

| **Dataset** | **Demixer** | | | **QuantTB** | | |
|---|---|---|---|---|---|---|
| | **Precision** | **Recall** | **$F1$ score** | **Precision** | **Recall** | **$F1$ score** |
| **10x** | 0.95 | 0.88 | 0.91 | 1.00 | 0.79 | 0.87 |
| **20x** | 0.97 | 0.91 | 0.93 | 1.00 | 0.81 | 0.88 |
| **90x-10x** | 0.96 | 0.96 | 0.95 | 1.00 | 0.82 | 0.88 |
| **70x-30x** | 0.95 | 0.96 | 0.95 | 1.00 | 0.82 | 0.88 |

### 4. Supplementary Figures

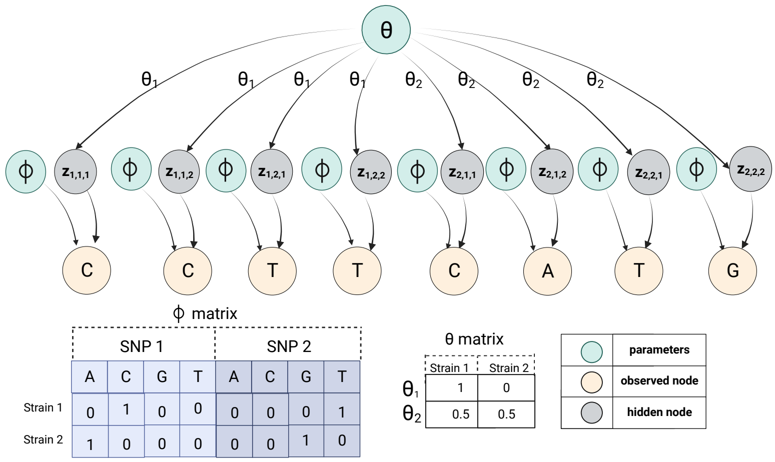


**Fig. S1. Generative process of SNP-LDA:** The assignment of strains to reads in a sample set consisting of 2 samples is shown. Each sample is assumed to have 2 SNPs with 2 reads per SNP. $\boldsymbol{\theta}$ and $\boldsymbol{\phi}$ are the model parameters and are assumed to be known.

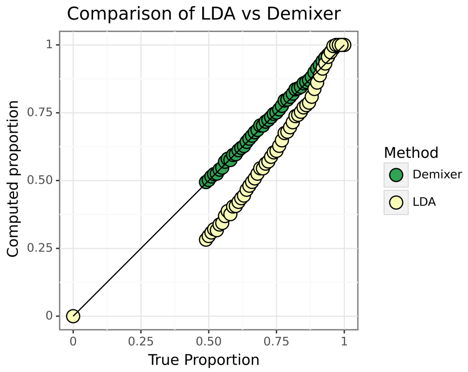


**Fig. S2. Comparison of traditional text LDA with Demixer:** The proportion of one of the strains estimated by traditional text LDA vs. Demixer on a 2-strain dataset (see Supplementary Section 1.6) is compared with its true proportion.

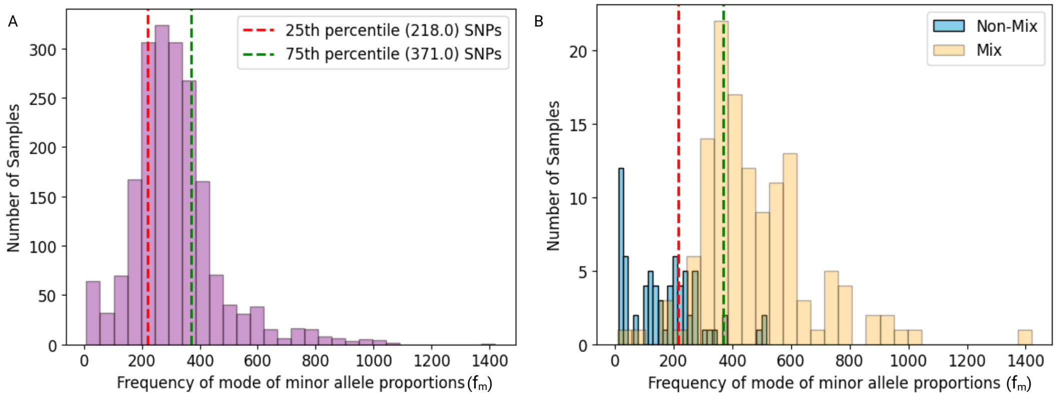


**Fig. S3. Distribution of $f_m$ values of Malawi samples:** The frequency of mode of minor allele proportions, denoted $f_m$ and expressed in terms of the number of SNPs, is used to assess the quality of mixed infection calls. A) depicts the 25th and 75th percentiles of $f_m$ values across all samples B) shows the histogram of $f_m$ of those samples called as mixed infection by Demixer but non-mixed by MixInfect (labelled as "Mix" and colored orange), and a second histogram corresponding to samples called as non-mixed by Demixer but mixed by MixInfect (labelled as "Non-mix" and colored blue); see also Figure 6C in main text.

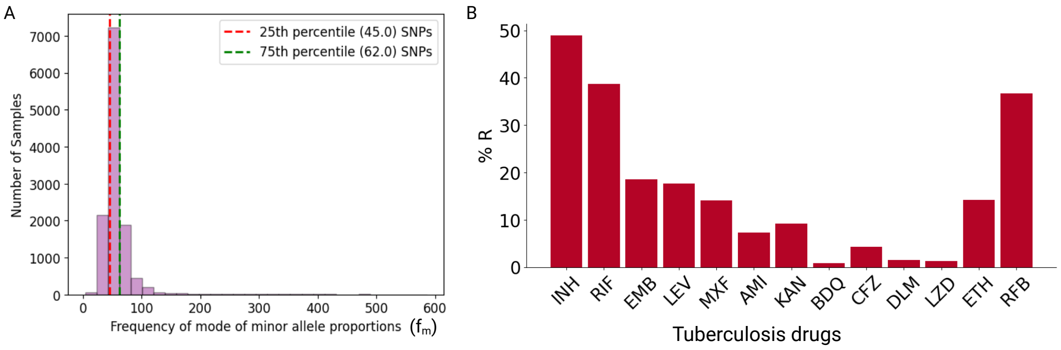


**Fig. S4. Analysis of CRyPTIC samples:** A) depicts the 25th and 75th percentiles of $f_m$ across all samples. The unit of $f_m$ is same as in previous figure. B) shows the percentage of mixed infection samples resistant to the 13 TB drugs isoniazid (INH), rifampicin (RIF), ethambutol (EMB), levofloxacin (LEV), moxifloxacin (MXF), amikacin (AMI), kanamycin (KAN), bedaquiline (BDQ), clofazimine (CFZ), delamanid (DLM), linezolid (LZD), ethionamide (ETH) and rifabutin (RFB). The figure B is generated using the opensource code from [4].

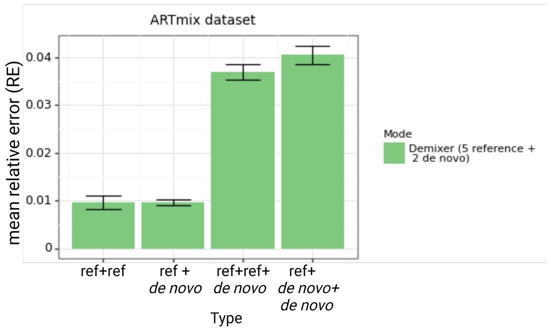


**Fig. S5. Demixer in *de novo* mode:** Demixer is ran on ARTmix dataset in *de novo* mode (see Section 3.3 in main text) with $\boldsymbol{wt}_{n,v}$ set to 1. The mean reative error is lower when compared to running Demixer with default weight 3 (refer Figure 5C).

#### 5. Supplementary Files

Supplementary data/result files listed below are available at this link:
https://drive.google.com/drive/folders/13WFACrn2EpeVTO7533-YwlAGjgF4UH3k?usp=drive_link.
Suppl File D1: The actual lineages of strains mixed and the lineages of the strains determined by Demixer and QuantTB for each sample in the *in vitro* dataset.
Suppl File D2: The strains, proportions, and confidence values inferred by Demixer for each sample in the Malawi dataset.
Suppl File D3: The strains and proportions estimated by Demixer for each sample in the CRyPTIC dataset. The susceptibility of samples to all 13 drugs provided as metadata in the CRyPTIC dataset is also included here for convenience.
Suppl File D4: The strains and proportions estimated by Demixer (trained using CRyPTIC dataset) for 10 samples in the Malawi dataset.
